# ROS signaling-induced mitochondrial Sgk1 regulates epithelial cell plasticity

**DOI:** 10.1101/2022.12.23.521432

**Authors:** Yingxiang Li, Chengdong Liu, Luke Rolling, Veronica Sikora, Zhimin Chen, Jack Gurwin, Caroline Barabell, Jiandie Lin, Cunming Duan

## Abstract

Many types of differentiated cells can reenter the cell cycle upon injury or stress. The mechanisms underlying this cell plasticity are still poorly understood. Here we investigated cell plasticity regulation using a zebrafish model, in which a population of differentiated epithelial cells are reactivated under a physiological context. We observed a robust and sustained increase in mitochondrial membrane potential in reactivated cells. Genetic and pharmacological perturbations show that elevated mitochondrial metabolism and ATP synthesis are critical for cell reactivation. Elevated mitochondrial metabolism increases mitochondrial ROS levels, which induces Sgk1 expression in the mitochondria. Deletion and inhibition of Sgk1 in zebrafish abolished cell reactivation. Similarly, ROS-dependent mitochondrial expression of SGK1 promotes S phase entry in human breast cancer cells. Mechanistically, Sgk1 coordinates mitochondrial activity with ATP synthesis by modulating F_1_F_o_-ATP synthase phosphorylation. These findings suggest a conserved intramitochondrial signaling loop regulating epithelial cell renewal.

**One sentence highlight:** This study reports a new intramitochondrial signaling loop regulating epithelial cell renewal.

## Introduction

Tissue homeostasis and regeneration require a tight control of cell renewal. Several adult tissues such as intestine, epidermis, and testis are endowed with robust cell renewal capabilities (Guillot and Lecuit, 2013; Tai et al., 2019). In mammalian small intestinal epithelium, for instance, cells are replaced every four to five days throughout life. This was initially attributed to a population of adult stem cells known as Lgr5+ crypt base columnar cells (CBCs) located in the base of the intestinal crypt (Clevers, 2013; Metcalfe et al., 2014). Subsequent studies revealed that other precursor cells in the crypt can revert to a Lgr5+ cell state to carry out the cell renewal function when the Lgr5+ CBS cells are genetically ablated (Clevers, 2013; Montgomery et al., 2011). More recent studies suggested that nearly all intestinal epithelial cells, including fully differentiated cells, have the ability to reenter the active cell cycle and replace the lost cells after injuries (e Melo and de Sauvage, 2019; Guiu et al., 2019; Post and Clevers, 2019). This phenomenon is not unique to the intestine or other tissues with rapid turnover rates, but also observed in tissues with very low physiological turnover rates such as heart, liver, lung, prostate etc. (Clevers and Watt, 2018). For example, upon partial hepatectomy, mammalian hepatocytes reenter the cell cycle and proliferate to regenerate the liver (Gadd et al., 2020). Another prominent example is the adult zebrafish heart, which displays robust regeneration after injuries (Poss et al., 2002). Decades of studies show that adult zebrafish hearts do not appear to have a stem cell pool. Rather, the newborn cardiomyocytes are derived from differentiated cardiomyocytes (Tzahor and Poss, 2017). These and other findings have led to the emerging notion that many types of differentiated cells are endowed with the ability to reenter the active cell cycle (e Melo and de Sauvage, 2019). Mounting evidence suggests that unlocking cell plasticity is a major hallmark of cancer cells (Hanahan, 2022). While it is becoming evident that this cell plasticity plays a key role in tissue homeostasis, regeneration, tumor formation and progression (e Melo and de Sauvage, 2019; Post and Clevers, 2019), the underlying mechanisms are still poorly understood. A major challenge in the field is that our current knowledge is mainly derived from injury models. Whether fully differentiated cells can de-differentiate and re-enter the cell cycle in a physiological context is still not clear (Clevers and Watt, 2018; Post and Clevers, 2019).

Recently, we have developed a zebrafish model, *Tg(igfbp5a:GFP)* fish (Dai et al., 2014; Liu et al., 2017). In this model, Ca^2+^-transporting epithelial cells, known as ionocytes or Na^+^-K^+^-ATPase-rich (NaR) cells, are genetically labeled by GFP expression (Dai et al., 2014; Liu et al., 2017). NaR cells are structurally and functionally similar to human intestinal and renal epithelial cells and contain similar molecular machinery for Ca^2+^ transcellular transport (Yan and Hwang, 2019). While distributed in the adult intestine, kidney, and gills, these polarized epithelial cells are located on the yolk sac epidermis during the larval stage (Hwang, 2009), making them easily accessible for experimental observation and manipulation. When zebrafish larvae are kept in an embryo medium containing normal concentration of [Ca^2+^] (0.2 mM, referred as the control medium), these polarized cells remain quiescent. When transferred to an embryo medium containing a low concentration of [Ca^2+^] (0.001 mM. referred as the induction medium hereafter), NaR cell number rapidly increases (Dai et al., 2014; Liu et al., 2017). BrdU pulse labeling, cell cycle analysis, and *in vivo* real-time cell tracing experiments have shown that the increased NaR cell number is due to the reactivation and proliferation of existing and differentiated NaR cells (Dai et al., 2014; Liu et al., 2017). Using this model, chemical biology screens and genetic studies have elucidated a number of cell autonomous and non-autonomous factors regulating NaR cell reactivation; they all converge onto the insulin-like growth factor (IGF)-PI3 kinase-Tor signaling pathway (Li et al., 2021; Liu et al., 2020; Liu et al., 2018; Xin et al., 2021; Xin et al., 2019). These findings are in good agreement with the roles of IGF/insulin-PI3 kinase-mTOR signaling in adult stem cell and T cell reactivation in mammals and *Drosophila* (Chell and Brand, 2010; Chen et al., 2008; Chen et al., 2009; Ziegler et al., 2019; Ziegler et al., 2015), suggesting a general and evolutionarily conserved mechanism at work.

The IGF/insulin-PI3 kinase-mTOR signaling is a nutrient sensitive and evolutionarily ancient pathway regulating metabolism and other biological processes (Mossmann et al., 2018). mTOR is a major regulator of mitochondrial activity and metabolism. Although mitochondrial metabolism has traditionally been viewed as a consequence of cellular state, recent studies suggest that mitochondrial metabolism plays key roles in dictating cell fate and cell state (Chakrabarty and Chandel, 2021; Nakamura-Ishizu et al., 2020; Rath et al., 2020). Two essential and interconnected functions of mitochondria are the tricarboxylic acid (TCA) cycle and oxidative phosphorylation (OXPHOS) (Chakrabarty and Chandel, 2021; Martínez-Reyes et al., 2016). The TCA cycle is critical in ATP production and in providing metabolic intermediates for biosynthesis and epigenetic regulation (Chakrabarty and Chandel, 2021). OXPHOS involves a directional flow of protons from the mitochondrial matrix through the ETC complexes into the intermembrane space, which forms an electrochemical gradient (Mitchell, 1966; Morelli et al., 2019). Mitochondrial membrane potential (Δ*Ψ*_m_) is a key indicator of the electrochemical gradient and mitochondrial activity (Zorova et al., 2018). Together with the proton gradient (ΔpH), ΔΨ_m_ forms the transmembrane potential, which drives ATP synthesis (Zorova et al., 2018). In addition to ATP production, OXPHOS also produces reactive oxygen species (ROS). Although initially viewed as mere by-products of mitochondrial metabolism, it is now understood that ROS also function as important signaling molecules (Chakrabarty and Chandel, 2021).

In this study, we tested the hypothesis that mitochondrial activity and metabolism play a key role in regulating NaR cell fate and renewal. We provided evidence that mitochondrial activity and metabolism are crucial in promoting NaR cell reactivation and proliferation. Mechanistically, elevated mitochondrial metabolism increases ATP synthesis and mitochondrial ROS production. ROS signaling induces the expression and mitochondrial localization of serum- and glucocorticoid-regulated kinase 1 (Sgk1), a member of the conserved AGC family protein kinases, which acts downstream in the PI3 signaling pathway. Sgk1 coordinates mitochondrial activity/metabolism change with ATP synthesis by modulating the F_1_F_o_-ATP synthase phosphorylation state. This signaling loop promotes both zebrafish NaR cell reactivation and human breast cancer cell S phase entry.

## Results

### Elevated mitochondrial activity acts downstream of IGF signaling to promote NaR cell reactivation

Using *Tg(igfbp5a:GFP)* larvae, in which GFP-labeled NaR cells located on the yolk sac surface and can be imaged directly in vivo (Dai et al., 2014; Liu et al., 2017), we measured mitochondrial membrane potential change (Δ*Ψ*_m_) by TMRM staining. Compared to quiescent, non-dividing NaR cells, a robust increase in Δ*Ψ*_m_ was detected in reactivated NaR cells (Figs. 1A and 1B). The elevation in Δ*Ψ*_m_ was detected in nearly all NaR cells within an hour after the induction and sustained for at least 48 hours (Fig. 1C). Similar results were obtained by MitoTracker-Red staining (Supplementary Figs. 1A-B). No change was detected in the mitochondrial mass, inferred by measuring the relative copy number of mitochondrial DNA to genomic DNA levels (Fig. 1D). Additionally, treatment with 2,4-Dinitrophenol (2,4-DNP) or Carbonyl cyanide 4-(trifluoromethoxy) phenylhydrazone (FCCP), two distinct uncouplers of mitochondrial oxidative phosphorylation, impaired NaR cell reactivation and proliferation (Figs. 1E-H). These results suggest that mitochondrial activity, but not mitochondrial mass, is elevated during NaR cell reactivation and that the elevated mitochondrial activity promotes cell reactivation. This idea was tested further by inhibiting the ETC complexes which mediate the directional proton flow (Mitchell, 1966; Morelli et al., 2019). Treatment of fish larvae with ETC complexes I, III, and IV inhibitors rotenone, antimycin, and sodium azide all inhibited NaR cell reactivation (Supplementary Fig. 2).

**Fig. 1.**
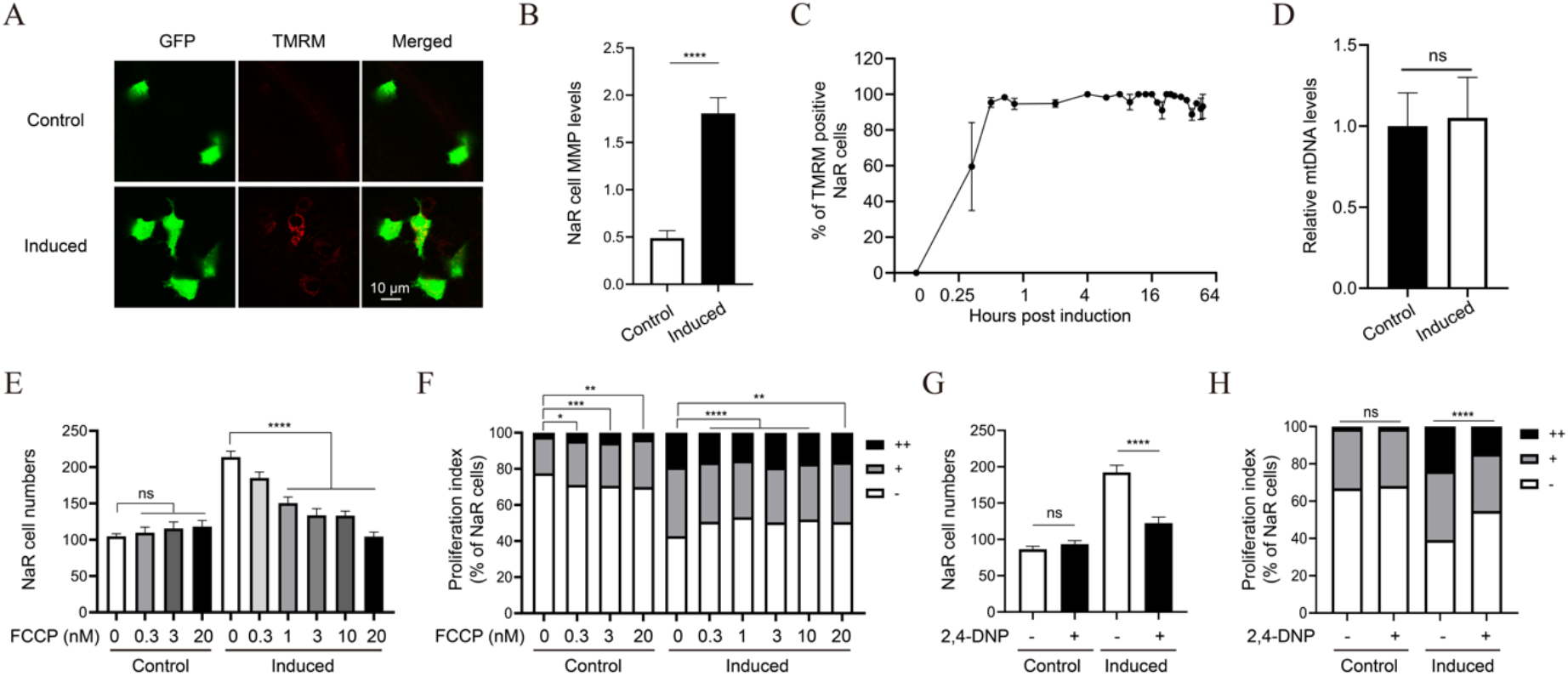
Elevated mitochondrial membrane potential (MMP) promotes NaR cell reactivation. (A-B) MMP levels in control and reactivated NaR cells. *Tg(igfbp5a:GFP)* larvae (3 dpf) were transferred to the control or induction medium. Two days later, MMP levels in NaR cells were measured after TMRM staining and normalized by GFP signal. Representative images are shown in (A) and quantified results in (B). n = 221~447 cells from multiple fish. (C) Time course of MMP change. *Tg(igfbp5a:GFP)* larvae (3 dpf) were transferred into the induction medium. A subset of larvae were randomly sampled at the indicated time point and % of TMRM-positive NaR cells were measured and shown. n = 2 ~ 6 fish/time point. (D) Mitochondrial DNA levels. NaR cells were isolated by FACS from fish described in (A-B). The levels of mitochondrial 16S rRNA gene were measured and normalized by the levels of nuclear aryl hydrocarbon receptor 2 gene. n = 4. (E-H) *Tg(igfbp5a:GFP)* larvae (3 dpf) were transferred to the control or induction medium containing the indicated concentration of phenylhydrazone (FCCP) (E and F) or 1μM 2,4-Dinitrophenol (2,4-DNP) (G and H). Two days later, NaR cell number (E, G) and NaR cell proliferation index (F, H) were determined and shown. n = 11~26 fish/group. The NaR cell proliferation index was determined by counting NaR cells that divided 0, 1, or 2 times (denoted by -, +, and ++) over the course of the experiment and presented as % of total NaR cells. In this and all subsequent figures, ns, not significant. *, **, *** and **** indicate *p* < 0.05, 0.01, 0.001, and 0.0001, respectively.

We next investigated the relationship between elevated Δ*Ψ*_m_ and IGF signaling. Previous studies have shown that NaR cell reactivation in *papp-aa*^-/-^ fish, which lacks the IGF binding protein 5 proteinase Papp-aa, is impaired due to defective IGF signaling in these cells (Liu et al., 2020). In contrast, NaR cells in *trpv6*^-/-^ fish, which are deficient in the epithelial calcium channel Trpv6, proliferate unrestrictedly due to constitutively elevated IGF signaling (Xin et al., 2019). If the elevated Δ*Ψ*_m_ in reactivated NaR cells is regulated by IGF signaling, then one would predict Δ*Ψ*_m_ in *pappaa*^-/-^ NaR cells would decrease. Likewise, elevated Δ*Ψ*_m_ should be detected in *trpv6*^-/-^ NaR cells. Indeed, while the induction medium treatment resulted in a robust increase in Δ*Ψ*_m_ in the NaR cells of wild-type and heterozygous sibling fish, this increase was not observed in the NaR cells of *pappaa*^-/-^ fish (Supplementary Fig.3A). In *trpv6*^-/-^ fish, NaR cells exhibited elevated Δ*Ψ*_m_ when kept in the control medium (Supplementary Fig.3B). To test this further, we treated *Tg(igfbp5a:GFP)* larvae with BMS-754807, an IGF1 receptor inhibitor (Carboni et al., 2009). Inhibition of the IGF1 receptor impaired NaR cell Δ*Ψ*_m_ increase in a dose dependent manner (Supplementary Fig. 3C). Finally, *Tg(igfbp5a:GFP)* fish were treated with rapamycin at a high centration that inhibits both Torc1 and Torc2 in zebrafish (Liu et al., 2017). Rapamycin treatment abolished the NaR cell Δ*Ψ*_m_ increase (Supplementary Fig. 3D). This is in agreement with previous studies showing BMS-754807 and rapamycin treatment abolished NaR cell reactivation (Liu et al., 2017). Collectively, these results suggest that elevated mitochondrial activity acts downstream of IGF signaling.

### Mitochondrial metabolism plays an essential role in NaR cell reactivation

The involvement of the TCA cycle/OXPHOS in NaR cell reactivation was investigated using zebrafish *noa*^-/-^ larvae, which lack pyruvate dehydrogenase (PDH) function due to the deletion of the PDH E2 subunit dihydrolipoamide s-acetyltransferase *(dlat)* (Taylor et al., 2004). While NaR cell reactivation was observed in the wild type and heterozygous siblings, it was impaired in the *noa*^-/-^ fish (Figs. 2A-B). Likewise, inhibition of PDH by CPI-613 (Zachar et al., 2011) (Supplemental Fig. 4A) reduced NaR cell reactivation in a dose-dependent manner (Figs. 2C-E). CPI-613 treatment lowered NaR cell proliferation in *trpv6^-/-^* fish to that of its siblings as well (Supplemental Figs. 4B-C). Inhibition of pyruvate transport from the cytoplasm into the mitochondria by α-cyano-4-hydroxycinnamate (CHC) (Del Prete et al., 2004) also impaired NaR cell reactivation (Supplemental Figs. 4D-E). Gboxin, an inhibitor of OXPHOS (Shi et al., 2019), had a similar effect (Figs. 2F-G).

**Fig. 2.**
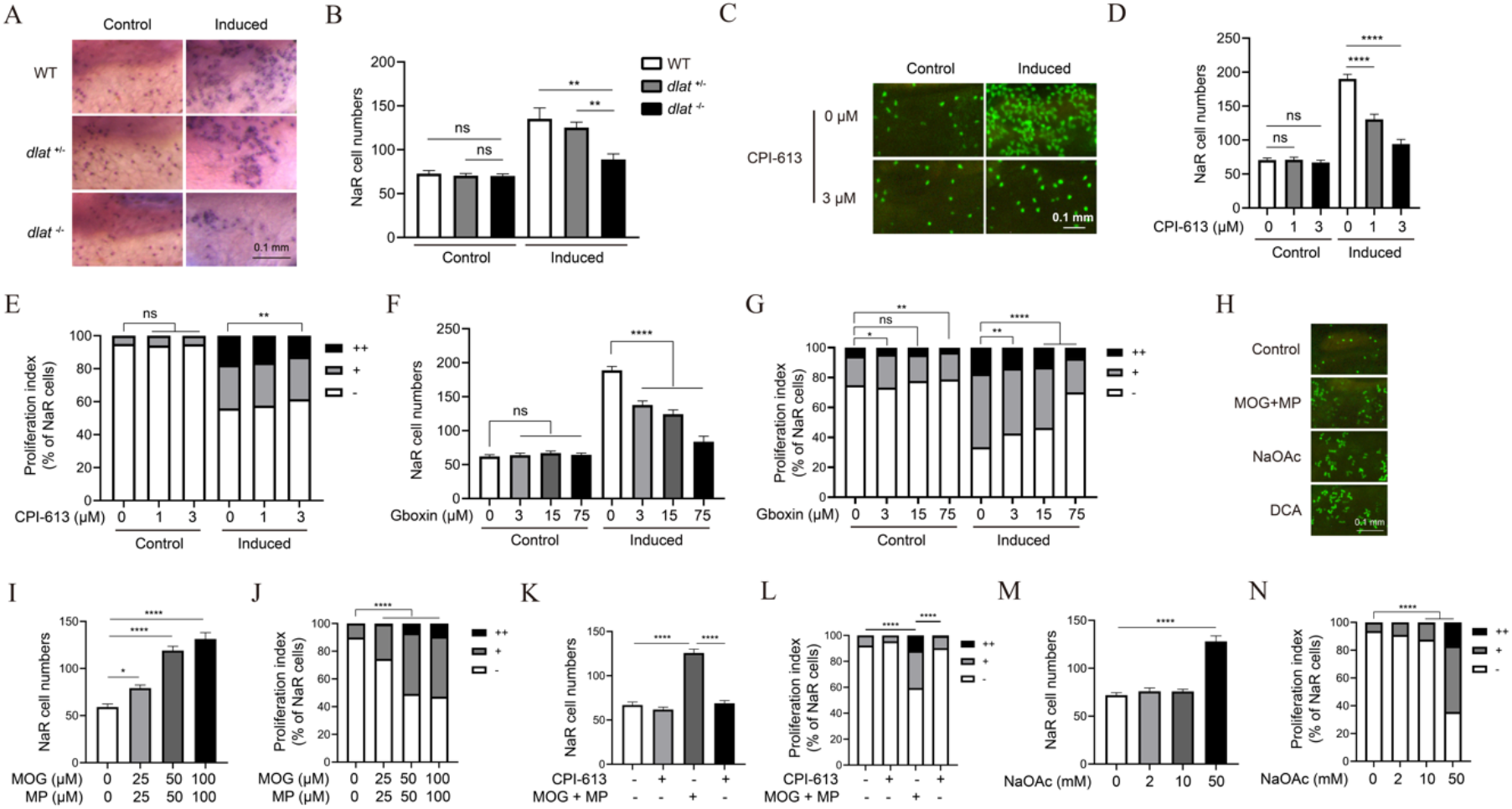
Elevated mitochondrial metabolism is required and sufficient to reactivate NaR cells. (A-B) Genetic disruption of the pyruvate dehydrogenase (PDH) complex impairs NaR cell reactivation. The progenies of *dlat^+/-^* intercrosses were transferred to the control or induction medium at 3 dpf. Two days later, NaR cells were labeled by *in situ* hybridization and quantified. Fish were genotyped individually afterwards. Representative images are shown in (A), and NaR cell number in (B). n = 13~40 fish/group. (C-E) Effect of PDH inhibitor CPI-613. *Tg(igfbp5a:GFP)* larvae (3 dpf) were transferred into the control or induction medium containing the indicated doses of CPI-613. Two days later, NaR cell number (D) and proliferation index (E) were measured and shown. Representative images are shown in (C). n=25~31 fish/group. (F-G) Effect of Gboxin. *Tg(igfbp5a:GFP)* embryos were raised and treated as described in (C) with the indicated doses of Gboxin. NaR cell number (F) and proliferation index (G) were measured and shown. n= 24-33 fish/group. (H-N) Effects of Dimethyl 2-oxoglutarate (MOG)+methyl pyruvate (MP), sodium acetate (NaOAc), and CPI-613 (3 μM). *Tg(igfbp5a:GFP)* larvae were transferred to the control medium containing the indicated chemicals at 3 dpf. Two days later, NaR cell number (I, K, M) and proliferation index (J, L, N) were measured and shown. n = 15~31 fish/group. Representative images are shown in (H).

To test whether an increase in TCA cycle/OXPHOS activity is sufficient to promote NaR cell reactivation, fish were treated with dimethyl α-ketoglutarate (MOG) and methyl pyruvate (MP). MOG and MP can be cleaved to generate α-ketoglutarate and pyruvate in cells, supplying key intermediates so as to increase TCA cycle and mitochondrial activity (Liang et al., 2020). MOG+MP treatment elevated NaR cell Δ*Ψ*_m_ and increased NaR cell reactivation under the control condition (Fig. 2H-J; Supplemental Fig. 1C). Co-treatment with CPI-613 abolished the MOG+MP-induced Δ*Ψ*_m_ increase and NaR cell reactivation (Fig. 2K-L; Supplemental Fig. 1C). Similarly, treatment with dichloroacetate (DCA), which indirectly activates PDH by inhibiting pyruvate dehydrogenase kinase (Stacpoole, 1989) increased NaR cell reactivation (Supplemental Figs. 4F-G). Finally, addition of sodium acetate, a precursor of acetyl-coA (Tiwari et al., 2020) into the control medium was sufficient to induce NaR cell reactivation (Figs. 2M-N). Taken together, these results suggest that elevated TCA cycle/ OXPHOS is both necessary and sufficient to promote NaR cell reactivation.

### ATP synthesis and ROS signaling are involved in cell reactivation

Consistent with the above results, in vivo measurement detected a 4-fold increase in mitochondrial ATP levels in reactivated NaR cells (Figs. 3A-B). MOG+MP treatment increased NaR cell mitochondrial ATP levels in fish kept in the control medium and this increase was abolished by CPI-613 co-treatment (Fig. 3C). In continuously dividing NaR cells in *trpv6^-/-^* larvae, the mitochondrial ATP levels were 8-fold greater than those in their siblings (Fig. 3D). Addition of the F_1_F_o_-ATP synthase inhibitor oligomycin reduced the ATP levels to nearly undetectable levels (Fig. 3D). Oligomycin treatment also inhibited NaR cell reactivation (Figs. 3E-F). The idea that elevated ATP synthesis is required for NaR cell reactivation was tested further using a F0 CRISPR/Cas9 gene deletion method (Kroll et al., 2021; Wu et al., 2018). Deletion of *atp5b,* which encodes the β subunit of F_1_F_o_-ATP synthase, abolished NaR cells reactivation under the induction medium and by MOG-MP treatment (Figs. 3G-J; Supplementary Fig. 5). One caveat of the above manipulations is that they affect ATP synthesis in all cell types. To determine the role of endogenous ATP synthesis in NaR cells, ATPase Inhibitory Factor 1 (IF1) was expressed in a subset of NaR cells *Tg(igfbp5a:GFP)* fish using a Tol2 transposon-mediated genetic mosaic assay (Liu et al., 2018). IF1 binds to and inhibits F_1_F_o_-ATP synthase in a pH-dependent manner (Cabezon et al., 2001). IF1^H49K^ is a pH-insensitive and constitutively active form of IF1 (Formentini et al., 2014). When expressed randomly in a subset of NaR cells, IF1^H49K^ inhibited NaR cell reactivation and proliferation in a cell autonomous fashion (Fig. 3K). Likewise, NaR cell-specific expression of another constitutively active IF1 mutant, hIF1^S39A^ (García-Bermúdez et al., 2015), also impaired NaR cell reactivation (Fig. 3L).

**Fig. 3.**
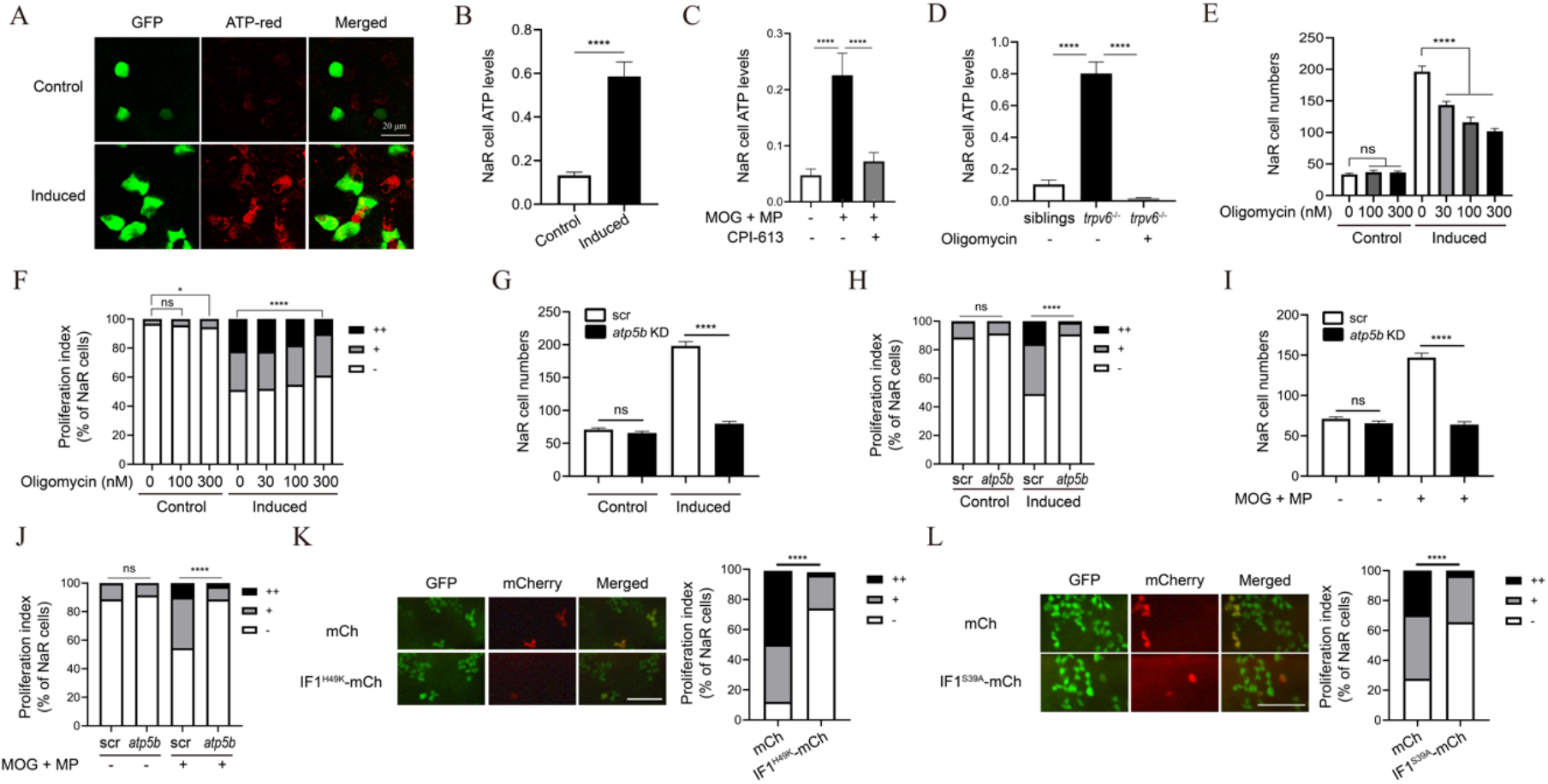
NaR cell reactivation requires ATP synthesis. (A-B) Elevated mitochondrial ATP levels in reactivated NaR cells. *Tg(igfbp5a:GFP)* larvae (3 dpf) were transferred to the control or induction medium. After 18 hours, NaR cell mitochondrial ATP levels were measured. Representative images are shown in (A) and quantified results in (B). n = 205~447 cells from multiple fish. (C) Effects of MOG + MP and CPI-613. *Tg(igfbp5a:GFP)* fish (3 dpf) were transferred to the control medium containing DMSO, 100 μM MOG + 100 μM MP, and/or 3 μM CPI-613. Five hours later, NaR cell mitochondrial ATP levels were measured and shown. n = 157~233 cells from multiple fish. (D) Elevated mitochondrial ATP levels in *trpv6^-/-^* NaR cells. Progeny from *trpv6^+/-^; Tg(igfbp5a:GFP)* intercrosses were raised to 3 dpf. They were transferred to the control medium with or without 200 nM Oligomycin. After measuring NaR cell mitochondrial ATP levels, the genotype of each fish was determined. n = 112 ~ 315 cells from multiple fish. (E-F) Effect of oligomycin. *Tg(igfbp5a:GFP)* fish (3 dpf) were transferred to the control or induction medium containing oligomycin at the indicated concentrations. Two days later, NaR cell number (E) and proliferation index (F) were measured and shown. n = 12 ~34 fish/group. (G-J) CRISPR-Cas9-mediated deletion of *atp5b. Tg(igfbp5a:GFP)* embryos injected with targeting gRNAs and *Cas9* mRNA were raised to 3 dpf and transferred into the control or induction medium without (G, H) or with 100 μM MOG + 100 μM MP (I-J). Two days later, NaR cell number (G, I) and proliferation index (H, J) were measured and shown. n = 14-42 fish/group. (K-L) NaR cell-specific expression of IF1^H49K^ (K) or IF1^S39A^ (L). *Tg(igfbp5a:GFP)* embryos injected with BAC-mCherry, BAC-IF1^H49K^ or BAC-IF1^S39A^-mCherry DNA were raised to 3 dpf and transferred into the induction medium. Two days later, NaR cells expressing the transgene were identified and cell proliferation index was measured. Representative images are shown in the left panel and quantified data in the right. Scale bar represents 0.1 mm. n = 102~163 in (K) and n = 29~148 cells from multiple fish in (L).

OXPHOS uses O_2_ as the final electron acceptor to produce ATP and this process also generates reactive oxygen species (ROS) (Rath et al., 2018; Shadel and Horvath, 2015). MitoSOX Red staining detected a significant increase in mitochondrial ROS levels (mtROS) in reactivated NaR cells (Figs. 4A-B). The signal is specific as it was abolished by the addition of an ROS scavenger, Mitoquinone (MitoQ) (Supplemental Fig. 6). The increase in mtROS was detected within 2 hours and sustained thereafter (Fig. 4C). As expected, MOG+MP treatment increased NaR cell mtROS levels (Fig. 4D) and CPI-613 co-treatment abolished this effect. The role of mtROS in NaR cell reactivation was tested using MitoQ, a mitochondrial ROS scavenger. MitoQ inhibited NaR cell reactivation in a dose-dependent manner (Figs. 4E-F). Conversely, treatment of fish with MitoParaquat (MitoPQ), which increases mtROS (Robb et al., 2015), induced NaR cell reactivation in a dose-dependent manner (Figs. 4G-H). These results suggest that mtROS signaling plays a crucial role in regulating NaR cell reactivation.

**Fig. 4.**
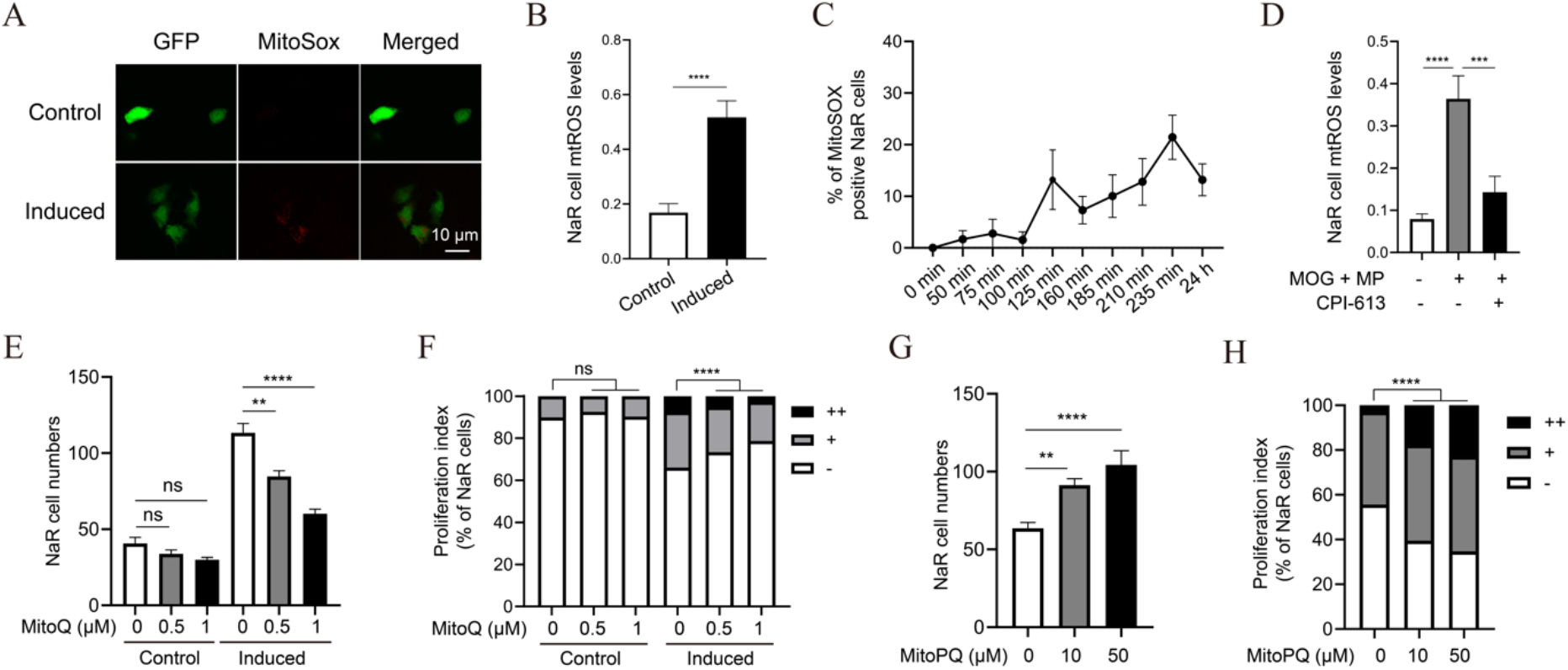
ROS signaling promotes NaR cell reactivation. (A-B) Mitochondrial reactive oxygen species (mtROS) levels. *Tg(igfbp5a:GFP)* larvae were transferred to the control or induction medium at 3 dpf. One day later, NaR cell mtROS levels were measured. Representative images are shown in (A) and quantified results in (B). n = 201~390 cells from multiple fish. (C) Time course changes. *Tg(igfbp5a:GFP)* larvae (3 dpf) were transferred into the induction medium. A subset of larvae were randomly sampled at the indicated time point and % of MitoSOX-positive NaR cells were determined and shown. n = 4 fish/group. (D) Effects of MOG+MP and CPI-613. *Tg(igfbp5a:GFP)* larvae (3 dpf) were transferred into the control or induction medium with or without 100 μM MOG + MP and 3 μM CPI-613. One day later, NaR cell mtROS levels were measured and shown. n= 197~276 cells from multiple fish. (E-F) mtROS signaling is required. *Tg(igfbp5a:GFP)* larvae were transferred to the control or induction medium containing the indicated dose of MitoQ at 3 dpf. Two days later, NaR cell number (E) and proliferation index (F) were measured and shown. n = 15~42 fish/group. (G-H) ROS signaling is sufficient. *Tg(igfbp5a:GFP)* larvae were raised in the control medium containing the indicated concentrations of MitoParaquat (MitoPQ) from 3 to 5 dpf. NaR cell number (G) and proliferation index (H) were measured and shown. n = 15~42 fish/group.

### Sgk1 coordinates ROS signaling with ATP synthesis by modulating F_1_F_o_-ATP synthase phosphorylation

We next tested the idea that Sgk1 acts as an effector molecule downstream of the ROS signaling. SGK1/Sgk1 is a kinase which acts downstream of the PI3 kinase signaling pathway (Di Cristofano, 2017). SGK1 is phosphorylated and activated by PDK1 and mTORC2 (Castel et al., 2016; Zhou et al., 2021). Upon activation, SGK1 phosphorylates TSC2 to activate mTORC1 (Castel et al., 2016). Previous studies demonstrated that ROS signaling increases SGK1 expression in human cell culture systems (Jiang et al., 2019; Kobayashi et al., 1999; O’Keeffe et al., 2013). In zebrafish, H_2_O_2_ treatment increased *sgk1* mRNA levels in a dose-dependent manner (Fig. 5A). To investigate the role of Sgk1 in cell reactivation, we measured *sgk1* mRNA levels in reactivated and quiescent NaR cells. A significant increase was detected in reactivated cells (Fig. 5B). Next, zebrafish larvae were treated with an SGK1 inhibitor, GSK650394. It inhibited NaR cell reactivation in a dose-dependent manner (Fig. 5C-D). EMD-638683, a distinct SGK1 inhibitor, had a similar effect (Figs. 5E-F). Furthermore, CRISPR/Cas9-mediated *sgk1* deletion impaired NaR cell reactivation (Fig. 5G-H; Supplementary Fig. 7). This action is specific to Sgk1 because CRISPR/Cas9-mediated deletion of *sgk2* had no such effect (Supplemental Figs. 8). Finally, SGK1^K127M^, a dominant-negative form of SGK1, was expressed in NaR cells specifically using the Tol2 transposon-mediated genetic mosaic assay. SGK1^K127M^ expression impaired the reactivation in a cell-autonomous fashion (Fig. 5I). Human SGK1 has a putative mitochondrial targeting sequence and was found to localize to the mitochondria in mouse mammary gland epithelial cells (O’Keeffe et al., 2013) and in human breast cancer cells under sorbitol induced osmotic stress (Supplementary Fig. 9). This sequence is conserved in zebrafish Sgk1 (Supplemental Figure 9) but absent in zebrafish Sgk2 (Supplementary Fig. 8). When SGK1 and F_1_F_o_-ATP synthase were incubated together in vitro, SGK1 was capable of increasing F_1_F_o_-ATP synthase phosphorylation (O’Keeffe et al., 2013). We postulated that ROS signaling-induced SGK1/Sgk1 expression/activity may increase ATP production by altering F_1_F_o_-ATP synthase phosphorylation state. This idea was tested with HEK 293 cells because of antibody availability and the high transfection efficacy. H_2_O_2_ treatment significantly increased the levels of phosphorylated ATP5B (Fig. 5J). This increase was abolished by GSK650394, suggesting this is a SGK1-dependent event (Fig. 5J). Likewise, expression of SGK1^S422D^, a constitutively active form of SGK1, increased the levels of phosphorylated ATP5B and this effect was abolished by GSK650394 (Fig. 5K). Consistent with these in vitro results, GSK650394 treatment abolished the increase in ATP levels in NaR cells under the induction condition (Fig. 5L). Collectively, these data suggest that elevated ROS signaling induces Sgk1 expression, which promotes cell reactivation by modulating F_1_F_o_-ATP synthase phosphorylation state and ATP synthesis.

**Fig. 5.**
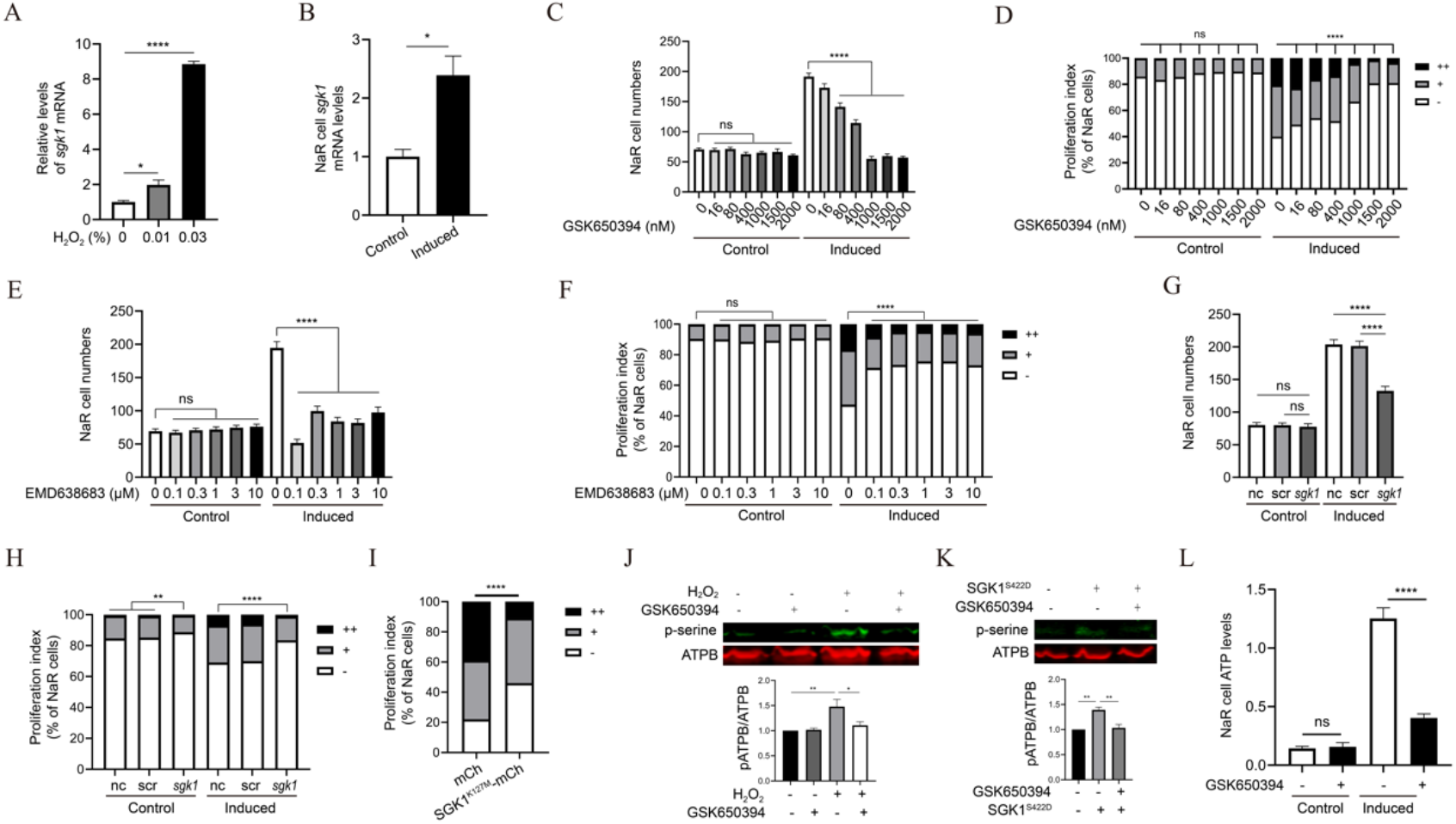
ROS signaling-induced Sgk1 expression is critical. (A) Induction of *sgk1* expression by H_2_O_2_. Zebrafish larvae(3 dpf) were transferred to the control medium containing the indicated doses of H_2_O_2_. After 3 hours, the larvae were collected and *sgk1* mRNA levels measured. n = 4. (B) *sgk1* mRNA levels in reactivated and quiescent NaR cells. NaR cells were isolated by FACS sorting from *Tg(igfbp5a:GFP)* larvae treated with the control or induction medium for 18 hours. *sgk1* mRNA levels were measured and shown. n=4. (C-F) Effects of SGK1 inhibitors. *Tg(igfbp5a:GFP)* embryos (3 dpf) were transferred to the control or induction medium containing the indicated doses of GSK-650394 (C-D) or EMD638683 (E-F). Two days later, NaR cell number (C and E) and proliferation index (D and F) were measured and shown. n = 15~41 fish/group. (G-H) CRISPR/Cas9-mediated *sgk1* deletion. *Tg(igfbp5a:GFP)* embryos were injected with *sgk1* targeting gRNAs and *Cas9* mRNA at the one-cell stage. They were raised to 3 dpf and transferred to the control or induction medium. Two days later, NaR cell number (G) and proliferation index (H) were measured and shown. n = 30~58 fish/group. (I) Effect of SGK1^K127M^ expression in NaR cells. *Tg(igfbp5a:GFP)* embryos were injected with the indicated BAC-mCherry DNA at the one-cell stage. They were raised to 3 dpf and transferred into the induction medium. Two days later, cell proliferation index in mCherry expressing NaR cells was determined and shown. n = 45~211 from multiple fish. (J) HEK293 cells were treated with DMSO or GSK650394 (20 μM) overnight, followed by 2 hour H_2_O_2_ (250 μM) treatment. Cells were subjected to IP using an anti-ATP5B antibody. The IP samples were analyzed by western blot using the indicated antibodies. Representative results are shown in the top panel. Ratio of Phospho- and total ATP5B is shown at the bottom. n =4. (K) HEK293 cells were transfected with Sgk1^S422D^-mcherry or an empty vector, followed by GSK650394 (20 μM) treatment. Cells were analyzed and shown as described in (J). n =3. (L) Elevated mitochondrial ATP synthesis in reactivated NaR cells is Sgk1-dependent. *Tg(igfbp5a:GFP)* larvae (3 dpf) were transferred to the control or induction medium for 14 hours. They were then treated with DMSO or GSK650394 (1 μM) for 4 hours. ATP levels were analyzed and shown. n = 178~609 cells from multiple fish.

Genetic studies in *C. elegans* suggest that SGK1 acts downstream of TORC2 to inhibit autophagy (Aspernig et al., 2019; Zhou et al., 2019). SGK1 has been shown to inhibit autophagy in cultured human cells (Zuleger et al., 2018). To determine whether autophagy is involved in NaR cell reactivation, zebrafish were treated with hydroxychloroquine (HCQ), which inhibits autophagy by blocking the fusion of autophagosomes with lysosomes (Solomon and Lee, 2009). HCQ treatment had little effect on NaR cell reactivation (Supplemental Figs. 10A-B). The involvement of autophagy was further investigated using the autophagy defective *gnptab^-/-^* zebrafish (Lu et al., 2020; Willet et al., 2018). No difference was detected in NaR cell reactivation between *gnptab^-/-^* fish and their siblings, indicating that autophagy does not play a major role in NaR cell reactivation (Supplemental Figs. 10C).

### SGK1 mediates ROS signaling-induced human breast cancer cell proliferation

We employed human MDA-MB-231 breast cancer cells to investigate the role of ROS-dependent mitochondrial expression of SGK1 in cell cycle regulation. Human MDA-MB-231 breast cancer cells undergo proliferation in response to increases in mitochondrial metabolism (Yang et al., 2018). H_2_O_2_ treatment of MDA-MB-231 cells strongly induced SGK1 mitochondrial localization (Supplementary Fig. 9B). MOG treatment resulted in a significant increase in mtROS levels (Fig. 6A). MOG treatment increased the percentage of cells entering the S phase and this effect was abolished by MitoQ (Fig. 6B). Conversely, treatment of cells with MitoPQ, which increases mitochondrial ROS (Fig. 6A), stimulated cells to enter S phase (Fig. 6C). Inhibition of SGK1 by GSK-650394 abolished both MOG- and MitoPQ-induced cell proliferation (Figs. 6D-E). Likewise, siRNA-mediated SGK1 knockdown inhibited cell proliferation in response to MOG and MitoPQ treatments (Figs. 6F-G). Next, seahorse assays were carried out to measure oxygen consumption rate (ORC) changes. As shown in Fig. 6H-I, MOG treatment increased the basal, the maximal mitochondrial respiratory capacity, and ATP production in MDA-MB-231 breast cancer cells. This effect was inhibited by GSK-650394 co-treatment (Fig. 6H-I). MOG treatment also increased proton leak and the spare or reserved capacity but GSK-650394 did not have significant effect on these parameters (Fig. 6H-I). Finally, inhibition of F_1_F_o_-ATP synthase by oligomycin treatment impaired MOG- and MitoPQ-stimulated cell proliferation (Figs. 6I-J). Taken together, these results suggest that SGK1 mediates mitochondrial metabolism and ROS signaling-induced breast cancer cell proliferation.

**Fig. 6.**
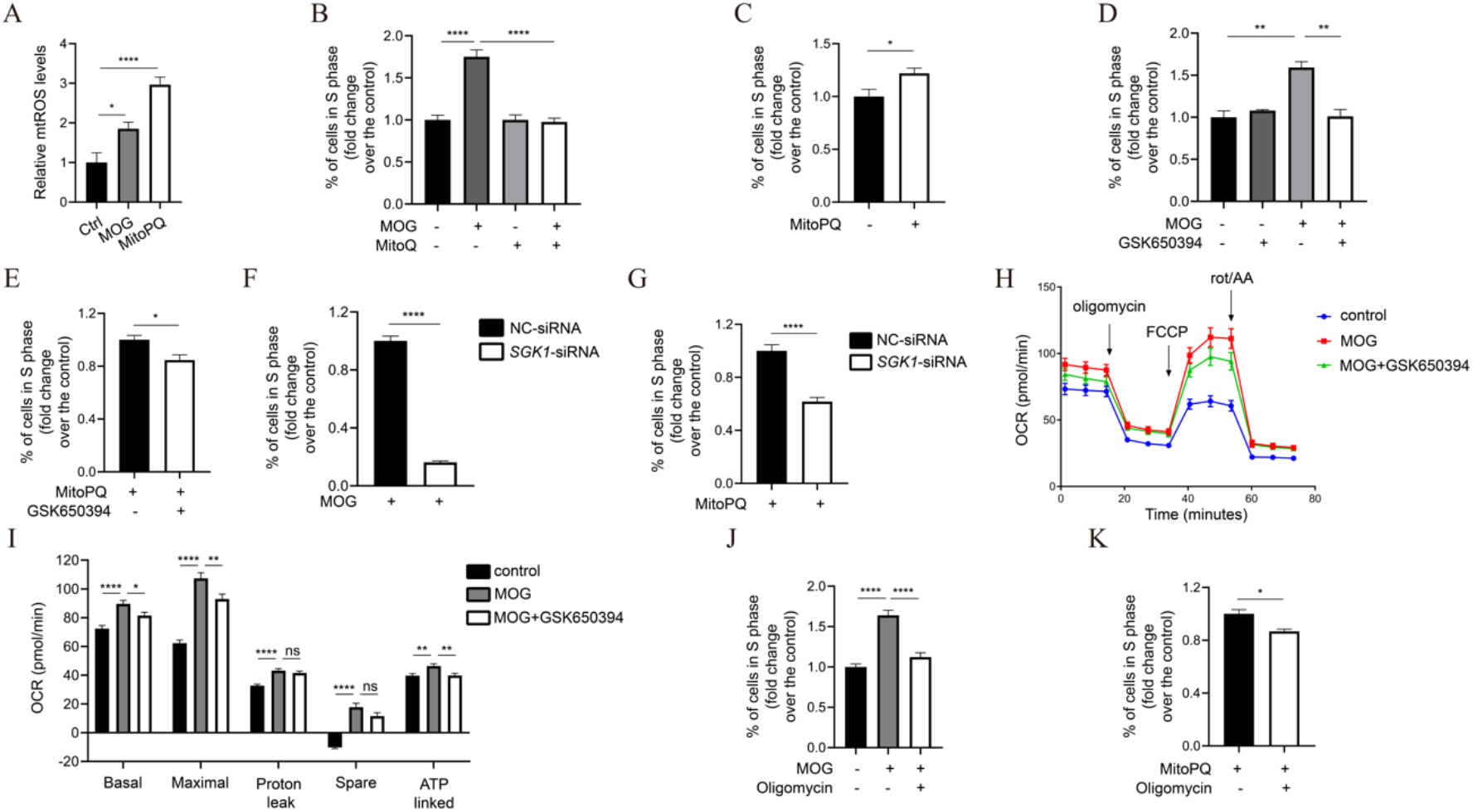
The mitochondrial metabolism-ROS-SGK1-ATP synthase signaling loop stimulates human breast cancer cell S phase entry. (A) MOG and MitoQ increase mtROS levels. MDA-MB-231 cells were treated with MOG (4 mM) or MitoPQ (5 μM) for 1 day. mtROS levels were measured and shown. n = 6. (B) mtROS signaling mediates MOG-induced cell proliferation. After MDA-MB-231 cells were treated with MOG (4 mM) and/or MitoQ (1 μM) for 2 days, % of cells in S phase was determined by flow cytometry and expressed as fold changes over the control group. n = 4. (C) Effect of MitoPQ. After MDA-MB-231 cells were treated with MitoPQ (5 μM) for 2 days, % of cell in S phase was determined and expressed as fold change over the control group. n = 7. (D-E) SGK1 mediates MOG- and MitoPQ-induced cell proliferation. MDA-MB-231 cells were treated with MOG (4 mM), MitoPQ (5 μM), and/or GSK650394 (20 μM) for 2 days. % of cells in S phase was determined and expressed as fold change over the control group. n = 3. (F-G) Knockdown of SGK1 abolishes MOG- (F) and MitoPQ (G)-induced cell proliferation. MDA-MB-231 cells were transfected with control (NC-siRNA) or *SGK1*-siRNA. One day after transfection, MOG (4 mM) or MitoPQ (5 μM) was added. % of cells in S phase was determined 2 days later and expressed as fold change over the control group. n = 6. (H-I) Inhibition of SGK1 reduces oxygen consumption rate. Mitochondrial oxygen consumption rate (OCR) in MDA-MB-231 cells treated with MOG and/or GSK650394 (20 μM) for 4 hours prior to seahorse assays. Oligomycin, FCCP and Rotenone/AA were spiked into cells at various timepoints. Oxygen consumption rate (OCR) was measured. Calculated parameters are shown in (I). n = 8. Similar results were obtained in a separate experiment. (J-K) F_1_F_o_-ATP synthase activity is required in MOG- (J) and ROS (K)-induced cell proliferation. After MDA-MB-231 cells were treated with MOG (4 mM), MitoPQ (5 μM) and/or Oligomycin (5 μM) for 2 days, % of cells in S phase was determined and expressed as fold change over the control group. n = 6.

## Discussion

In this study, we provided several lines of evidence suggesting that mitochondrial metabolism plays a key role in regulating epithelial cell fate and renewal. We found that mitochondrial activity and the TCA cycle/OXPHOS are maintained at relatively low levels in quiescent NaR cells. During cell reactivation, the mitochondrial activity and the TCA cycle/OXPHOS are elevated. This leads to elevated ATP synthesis and higher mtROS levels. Elevated mtROS signaling induces the expression and mitochondrial localization of Sgk1. Sgk1 links changes in mitochondrial metabolism/activity to ATP synthesis by modulating the F_1_F_o_-ATP synthase phosphorylation state. The ROS signaling-SGK1-F_1_F_o_-ATP synthase signaling loop not only plays a critical role in zebrafish NaR cell reactivation but also mediates mitochondrial metabolism-induced human breast cancer cell proliferation.

Zebrafish use NaR cells to take up Ca^2+^ from the surrounding aquatic habitat. When facing a low [Ca^2+^] environment, NaR cells are reactivated and undergo robust proliferation to yield more NaR cells in order to maintain body Ca^2+^ homeostasis (Dai et al., 2014; Hwang, 2009; Liu et al., 2017; Liu et al., 2018). Previous studies have shown that this adaptive reactivation of NaR cells is regulated by the IGF-PI3 kinase-Tor signaling activity (Dai et al., 2014; Liu et al., 2017; Liu et al., 2020; Liu et al., 2018; Xin et al., 2019). In the present study, we show that NaR cell reactivation is accompanied by a robust and sustained increase in Δ*Ψ*_m_. In vivo measurement detected a major increase in mitochondrial ATP levels in reactivated NaR cells. Given that we observed no significant changes in mitochondrial mass, these results indicate that the mitochondrial activity is elevated in reactivated cells. This increase is likely resulted from the activation of IGF-PI3 kinase-Tor signaling. This conclusion is supported by the Δ*Ψ*_m_ changes observed in *pappaa*^-/-^ and *trpv6*^-/-^ fish and by the fact that the Δ*Ψ*_m_ increase was abolished by inhibition of the IGF1 receptor and Toc1/2. It is also in good agreement with previous studies showing IGF and Tor1/2 signaling as essential in NaR cell reactivation (Dai et al., 2014; Liu et al., 2017; Liu et al., 2020; Liu et al., 2018; Xin et al., 2019). We provided several lines of independent evidence showing that the elevated mitochondrial activity and metabolism are critical in NaR cell reactivation. Dissipating the mitochondrial membrane potential by 2,4-DNP and FCCP impaired NaR cell reactivation. Treatment with inhibitors for ETC complexes I, III, and IV also impaired NaR cell reactivation. Genetic deletion and pharmacological inhibition of PDH abolished NaR cell reactivation. Conversely, increasing TCA cycle/OXPHOS activity by MOG+MP treatment, sodium acetate, and other TCA cycle activators were sufficient to promote NaR cell reactivation. These results argue strongly that mitochondrial metabolism plays a key role in NaR cell reactivation.

Most published studies on the role of mitochondrial metabolism in epithelial renewal focus on adult stem cells. Whether these genetic manipulations alter the state of differentiated epithelial cells has not been fully explored. Perekatt and colleagues (Perekatt et al., 2014) reported that knockout of the transcriptional repressor YinYang 1 (YY1) in mouse intestinal stem cells up-regulated ETC genes and increased OXPHOS. This resulted in stem cell reactivation, proliferation, and eventually exhaustion (Perekatt et al., 2014). Another study showed that deletion of the glycolytic enzyme pyruvate kinase M2 isoform (Pkm2) in mice resulted in elevated OXPHOS and ATP production in intestinal stem cells, and this led to the development of colon cancer (Kim et al., 2019). Similarly, intestinal stem cells in mutant *Drosophila* carrying a mutation impairing ETC proliferated at a slower rate (Zhang et al., 2020). These findings imply that an increase in OXPHOS is necessary to reactivate quiescent intestinal stem cells and promote their proliferation. In contrast, Rodríguez-Colman et al. reported that quiescent intestinal stem cells display high mitochondrial activity and greater OXPHOS compared with Paneth cells (Rodríguez-Colman et al., 2017). Inhibition of mitochondrial activity impairs intestinal stem cell reactivation (Rodríguez-Colman et al., 2017). Along the same line, genetic deletion of the mitochondrial pyruvate carrier (MPC) in mouse intestinal stem cells increased their proliferation rate, while MPC overexpression suppressed it (Schell et al., 2017). The reasons underlying these conflicting findings are not entirely clear, but these studies all used injury assays in stable mutant animals that are permanently deficient in one or more genes. Recent studies suggest that permanent knockouts can cause transcriptional adaptation and/or other compensatory mechanisms due to the inherent genetic robustness in multicellular organisms (Jakutis and Stainier, 2021). Our model differs from these studies in that we investigated the reactivation of a population of differentiated epithelial cells under a physiological stress. We used a combination of permanent and transient gene deletion, pharmacological inhibition/activation, and genetic mosaic assays.

A key finding made in this study is that increased mtROS levels resulting from elevated mitochondrial metabolism promotes cell reactivation by inducing Sgk1 expression in the mitochondria. Several lines of evidence support this conclusion. First, there is a significant increase in mtROS levels in reactivated NaR cells. The elevated mtROS are functionally important as NaR cell reactivation was abolished by treatment with a mtROS scavenger. Second, ROS treatment increased *sgk1* mRNA expression in zebrafish larvae in a dose-dependent manner. Third, ROS treatment resulted in a robust increase of SGK1 in the mitochondria in cultured human breast cancer cells. Inhibition of Sgk1/SGK1 by two distinct inhibitors impaired NaR cell reactivation in vivo and inhibited mitochondrial metabolism- and ROS-induced human breast cancer proliferation in vitro. Furthermore, CRISPR/Cas9-mediated deletion of *sgk1* blocked NaR cell reactivation. Likewise, siRNA-mediated knockdown of SGK1 inhibited breast cancer proliferation. Finally, dominant-negative inhibition of Sgk1 in NaR cells specifically impaired NaR cell reactivation. We postulate that Sgk1/SGK1 regulates cell reactivation by modulating F_1_F_o_-ATP synthase phosphorylation state and ATP synthesis. In support of this idea, a robust increase in mitochondrial ATP levels was detected in reactivated NaR cells. Similarly, there was a robust increase of mitochondrial ATP levels in continuously proliferating NaR cells in *trpv6^-/-^* mutant fish. Importantly, genetic deletion and pharmacological inhibition of Sgk1 reduced mitochondrial ATP levels and blocked NaR cell reactivation. Likewise, inhibition or siRNA-mediated knockdown of SGK1 abolished mitochondrial metabolism- and ROS-stimulated S phase entry of human breast cancer cells. Finally, inhibition of F_1_F_o_-ATP synthase by oligomycin treatment impaired NaR cell reactivation in vivo and human breast cancer cell proliferation in vitro. These findings are consistent with a recent report showing that SGK1 activity is both required and sufficient to increase ATP production in extracellular matrix detached (but not attached) human cancer cells, although increased GLU1-mediated glucose uptake was proposed to be the underlying mechanism (Mason et al., 2021). Although SGK1 is structurally related to AKT, it lacks the PH domain. A study reports that SGK1 activation occurs at a distinct subcellular compartment from that of Akt (Gleason et al., 2019). In this study, we found that SGK1/Sgk1 has a conserved mitochondrial targeting sequence and mitochondrial localization of SGK1 increases under elevated mitochondrial activity and ROS signaling. This is in agreement with a study reporting an increase of SGK1 expression in the mitochondria under osmotic stress (O’Keeffe et al., 2013). We presented biochemical evidence that ROS signaling increased ATP levels and ATP5B phosphorylation and that these increases were abolished by inhibiting SGK1/Sgk1 in vitro and in vivo. Moreover, expression of a constitutively active form of SGK1 was sufficient to increase the levels of phosphorylated ATP5B and this effect was abolished by a SGK1 inhibitor. These data are consistent with the notion that SGK1 likely affects ATP synthesis by modulating ATP5B phosphorylation state. Several F_1_F_o_-ATP synthase subunits are known to be post-translationally modified by protein kinases and these post-translational modifications are thought to be an important means of regulation of ATP synthesis (Nesci et al., 2017). Our sequence analysis reveals several putative SGK1 phosphorylation sites in human ATP5B, including T21, S23, S327, T348, T453, and S465. These sites are conserved in zebrafish Atp5b (T10, S49, S316, T337, T442, and S454). Future biochemical studies are needed to determine whether one or more of these sites are functional SGK1 phosphorylation site(s).

SGK1 has been shown to inhibit autophagy in cultured human cells (Zuleger et al., 2018). In *C. elegans, sgk1^-/-^* mutants exhibited elevated autophagy and increased mitochondrial permeability (mPTP) (Aspernig et al., 2019; Heimbucher et al., 2020). It was found that SGK1 phosphorylates VDAC-1, a mitochondrial outer membrane protein and a component of mitochondrial permeability (mPTP), and negatively regulates VDAC-1 by increasing its degradation (Zhou et al., 2019). Loss of MTORC2/SGK1 leads to an uncontrolled opening of the mPTP pore and increases autophagy (Zhou et al., 2019). Our pharmacological inhibition and genetic experiment results do not support a major role of autophagy in NaR cell reactivation. Interestingly, the *sgk1-/-* worms exhibited elevated mtROS levels but greatly reduced Δ*Ψ*_m_ (Aspernig et al., 2019; Heimbucher et al., 2020), suggesting SGK1 deficiency leads to defective mitochondrial homeostasis. These observations made in C. elegans, however, are different from studies reporting increased ΔΨ_m_ in mTORC2-deficient human cancer cells and mouse keratinocytes (Betz et al., 2013; Heimbucher et al., 2020; Tassone et al., 2017). The reasons for these different findings are not clear but it is plausible that there may be a feedback loop between mtROS- signaling and mitochondrial SGK1 expression/action and that modest increase in mtROS production may induce SGK1 expression but overproduction of mtROS suppress SGK1 expression.

In summary, the results of this study indicate a previously unrecognized intramitochondrial signaling loop which couples mitochondrial activity to ATP production and regulates epithelial cell plasticity. A key module in this regulatory loop is Sgk1 mitochondrial expression induced by ROS signaling. Mitochondrial Sgk1 changes cell state by stimulating ATP synthesis via modulating F_1_F_o_-ATP synthase phosphorylation, which in turn stimulates ATP synthesis. Given that approximately 90% of human cancers arise from epithelial cells and cell plasticity is a major hallmark of cancer (De Francesco et al., 2018), the regulation of epithelial cell plasticity by Sgk1 may have broader implications. Indeed, our results suggested a critical role of SGK1 in mediating mitochondrial metabolism- and ROS signaling-regulated S phase entry of human breast cancer cells. SGK1 was first discovered in rat tumor cells as a gene regulated by serum and glucocorticoids and has been implicated in ion channels and kidney function (Cicenas et al., 2022; Firestone et al., 2003; Webster et al., 1993). SGK1 was subsequently found to play a critical role in proliferation of cancer cells harboring PI3 kinase mutations (Castel et al., 2016; Orlacchio et al., 2017). Knockdown of SGK1 impaired basilar smooth muscle cell progress from G0/G1 phase to S phase (Chen et al., 2021) and overexpression of *SGK1* stimulates cultured colon cancer cell proliferation (Liang et al., 2017). SGK1 expression is elevated in colon cancer cells and in other cancers (Lang et al., 2010). *Sgk1-/-* mutant mice had reduced colon tumor formation when exposed to carcinogens (Nasir et al., 2009). Future studies are needed to determine the potential involvement of the ROS signaling-SGK1-F_1_F_o_-ATP synthase signaling loop in colon cancer and other epithelial derived cancers.

## Materials and methods

### Chemicals and Reagents

All chemical reagents were purchased from Fisher Scientific (Pittsburgh, PA) unless stated otherwise. Liberase, FCCP, 2,4-DNP, N-Acetyl-L-cysteine (NAC), α-Cyano-4-hydroxycinnamic acid (CHC), Dimethyl 2-oxoglutarate (MOG), Methyl pyruvate (MP), CPI-613, Sodium dichloroacetate (DCA), Sodium Acetate, Rotenone, Antimycin A, Sodium Azide, and Oligomycin were purchased from Sigma-Aldrich (St Louis, MO). Gboxin was purchased from Cayman Chemical (Ann Arbor, MI), Mitoquinone (MitoQ) from BioVision (Milpitas, CA), GSK-650394 from MedChem Express (Monmouth Junction, NJ), and EMD638683 from ApexBio (Houston, TX). Rapamycin was purchased from Calbiochem (Gibbstown, NJ). BMS-754807 was purchased from JiHe Pharmaceutica (Beijing, China). Cell culture media, fetal bovine serum, and Lipofectamine 3000 were purchased from (Invitrogen, Carlsbad, CA). The anti-ATPB and anti-β-actin antibodies were obtained from Proteintech (Rosemont, IL). The anti-phosphoserine antibody was obtained from Sigma-Aldrich (St Louis, MO) and anti-SGK1 antibody from Abcam (Cambridge, MA). Light chain specific anti-rabbit secondary antibody was purchased from Jackson ImmunoResearch (West Grove, PA). TRIzol LS reagent was purchased from (Invitrogen, Carlsbad, CA). Other secondary antibodies were purchased from Licor (Lincoln, NE). *Silencer* Select Negative Control No.1 siRNA was purchased from Fisher Scientific (Pittsburgh, PA) and MISSION esiRNA (targeting *SGK1*) was purchased from Sigma-Aldrich (St Louis, MO).

### Animals

Zebrafish were raised following standard guidelines (Westerfield, 2000). In addition to wild-type fish, *Tg(igfbp5a:GFP)* fish (Liu et al., 2017), *trpv6^+/-^; Tg(igfbp5a:GFP)* fish (Xin et al., 2019), and *pappaa^+/-^; Tg(igfbp5a:GFP)* fish (Liu et al., 2020) were used. The *noa* mutant line, *dlat*^-/-^ (Taylor et al., 2004) was obtained from the Zebrafish International Resource Center (ZIRC). Mutant *gnptab^-/-^* fish were obtained from Dr. Heather Flanagan-Steet, Clemson University (Lu et al., 2020). The fish were genotyped as previously reported (Liu et al., 2020; Xin et al., 2019). Embryos were obtained by natural crossing and staged following published protocols (Kimmel et al., 1995). All embryos were raised in the standard E3 embryo medium (Westerfield, 2000) until 3 dpf. In addition to E3 embryo medium, two additional media were used. The induction embryo medium containing 0.001 mM [Ca^2+^] and the control embryo medium containing 0.2 mM [Ca^2+^] were prepared following a previously reported formula (Dai et al., 2014). To prevent pigmentation, 0.003% (w/v) N-phenylthiourea (PTU) was added to the media when needed. All experiments were conducted in accordance with the guidelines approved by the University of Michigan Institutional Committee on the Use and Care of Animals.

### Mitochondrial membrane potential (MMP, Δ*Ψ*_m_) measurement

Δ*Ψ*_m_ was measured using tetramethylrhodamine (TMRM) or Mito-Tracker Red (Invitrogen, Carlsbad, CA). For TMRM staining, larval fish were incubated in the appropriate embryo medium containing 60 nM TMRM at 28.5 °C for 30 min. For MitoTracker-Red staining, *Tg(igfbp5a:GFP)* larvae were incubated in appropriate medium containing 150 nM MitoTracker-Red at 28.5 °C for 15 min. The fish were washed three times and imaged using a stereomicroscope (Leica MZ16F, Leica, Wetzlar, Germany) equipped with a QImaging QICAM camera (QImaging, Surrey, BC, Canada). Fluorescence intensity in each NaR cell was measured with Image J. Δ*Ψ*_m_ was evaluated by subtracting the background fluorescence intensity in NaR cells and normalized by GFP signal intensity.

### Fluorescence-activated cell sorting (FACS) and quantitative real-time PCR (qPCR) analysis

Isolation and sorting of NaR cells were performed as previously described (Liu et al., 2017). The sorted cells were collected and directly immersed in TRIzol LS reagent (Invitrogen, Carlsbad, CA) and kept at 4°C. RNA was isolated and cDNA was synthesized using SuperScript III Reverse Transcriptase. qPCR was performed on an ABI 7500 fast Real-Time PCR system (Applied Biosystems, Foster City, CA) using SYBR Green (Bio-Rad).

### Mitochondrial DNA measurement

Mitochondrial DNA levels were determined by measuring the mitochondrial 16s rRNA gene and normalized by a nuclear gene (aryl hydrocarbon receptor 2) following a published method (Hunter et al., 2010). Briefly, genomic DNA was extracted from FACS sorted NaR cells by digestion in a SZL buffer containing proteinase K (100 g/ml) for 2 hours at 60°C. The digestion was stopped by 15 min heat inactivation at 95°C. After centrifugation at 10,000 rpm for 1 min, the supernatant was subjected to qPCR to amplify a fragment of the mitochondrial 16S rRNA gene and a fragment of the aryl hydrocarbon receptor 2 gene (Hunter et al., 2010).

### Drug treatment

N-Acetyl-L-cysteine (NAC), Sodium Acetate, Sodium Azide, and DCA were first dissolved in water and then diluted in appropriate embryo media. Other drugs were first dissolved in DMSO first and then diluted in embryo media. *Tg(igfbp5a:GFP)* larvae were placed into a 6-well plate and washed three times with either the control or the induction medium or before the drug was added. The samples were collected and analyzed at the indicated time.

### Whole-mount *in situ* hybridization

Zebrafish larvae were fixed in 4% paraformaldehyde and permeabilized in methanol before use in whole-mount *in situ* hybridization, as described previously (Dai et al., 2014).

### In vivo measurement of mitochondrial ATP and ROS (mtROS) levels

NaR cell ATP and mtROS levels were measured with ATP-Red staining and MitoSOX staining, respectively. For this, *Tg(igfbp5a:GFP)* larvae were placed in appropriate embryo media containing 70 μM ATP-Red or 5 μM MitoSOX and incubated at 28.5 °C for ~15-20 min. After washing three times, the fish were imaged under a fluorescence microscope (Leica MZ16F, Leica, Wetzlar, Germany) equipped with a QImaging QICAM camera (QImaging, Surrey, BC, Canada) or a Leica TCS SP8 confocal microscope. Fluorescence intensity in each NaR cell was measured using ImageJ. ATP or mtROS levels were calculated by subtracting the background signal and normalized by GFP signal intensity in the same NaR cell.

### CRISPR/Cas9-mediated gene knockout

Cas9 mRNA was synthesized by *in vitro* transcription using the pT3.Cas9-UTRglobin plasmid as the template. Four sgRNAs were used to target one gene. The sequences were designed using CHOPCHOP (Labun et al., 2019). The primers used to synthesize gRNAs are shown in Supplementary Table 1. The sgRNAs were synthesized by *in vitro* transcription following a published method (Xin and Duan, 2018). Once synthesized, the sgRNAs (40 ng/μl) were mixed with Cas9 mRNA (400 ng/μl) and co-injected into *Tg(igfbp5a:GFP)* embryos at the one-cell stage. The injected embryos were raised in E3 embryo medium until 3 dpf and then transferred to the induction or control medium. Knockout efficacy was validated using headloop PCR following a published method (Kroll et al., 2021) and the primers are shown in Supplementary Table 1.

### Plasmid and BAC constructs

The ORF of human IF1 (IF1 ^H49K^) was sub-cloned into pIRES-mCherry using primers IF1-EcoRI-F and IF1-BamHI-R (Supplementary Table 1). IF1^S39A^ was engineered by two rounds of site-directed mutagenesis using IF1^H49K^ as a template. The primers used are *IF1SDM-S39AF, IF1SDM-S39AR, IF1SDM-K49HF and IF1SDM-K49HR* (Supplementary Table 1). The mouse Sgk1 ORF DNA was sub-cloned into pIRES-mCherry using primers Sgk1-EcoRI-F and Sgk1-BamHI-R (Supplementary Table 1). IF1^S39A^, IF1^H49K^ and Sgk1^K127M^ tagged with mCherry were then integrated to the *igfbp5aBAC* construct by replacing the *igfbp5a* coding sequence from the start codon to the end of the first exon via homologous recombination as previously reported (Liu et al., 2017). The primers used were igfbp5a-*IF1*-F, igfbp5a-*Sgk1*-F, and igfbp5a-pEGFP-C3-Kan-R (Supplementary Table 1). The resulted *BAC(igfbp5a:hIF1H49K-mCherry), BAC(igfbp5a:hIF1S39A-mCherry), BAC(igfbp5a:mSGK1K127M-mCherry)* constructs were confirmed by Sanger sequencing.

### Tol2 transposon-mediated mosaic transgenesis assay

A recently developed Tol2 transposon-mediated genetic mosaic assay was used to target transgene expression in a subset of NaR cells (Liu et al., 2018). For this, *BAC(igfbp5a:hIF1H49K-mCherry)*, *BAC(igfbp5a:hIF1S39A-mCherry), BAC(igfbp5a:mSGK1K127M-mCherry)* or *BAC(igfbp5a:mCherry)* DNA and Tol2 mRNA were mixed and injected into 1-cell stage *Tg(igfbp5a:GFP)* embryos. The embryos were raised and subjected to various treatments. Cells co-expressing mCherry and GFP were identified and the cell proliferation index was determined as previously reported (Liu et al., 2018).

### Cell cultures and transfection

Human embryonic kidney cells (HEK293) were purchased from the American Type Tissue Collection (ATCC). Human MDA-MB-231 breast cancer cells were obtained from Dr. Yanzhuang Wang’s lab at University of Michigan. HEK293 cells were cultured in DMEM/F-12 cell culture medium (Gibco, Carlsbad, CA) supplemented with 10% FBS, penicillin and streptomycin. MDA-MB-231 cells were cultured in DMEM cell culture medium (Gibco, Carlsbad, CA) supplemented with 5% FBS, penicillin, and streptomycin. Cells were maintained at 37 °C in a humidified atmosphere of 5% CO_2_. Plasmid transient transfection was performed using Lipofectamine 3000 (Invitrogen, Carlsbad, CA) and siRNA transfection was conducted using Lipofectamine RNAiMAX (Invitrogen, Carlsbad, CA) following the manufacturer’s instructions.

### Immunoprecipitation (IP)

Cultured cells were lysed using the RIPA buffer (Sigma-Aldrich, St Louis, MO). IP was performed with the Dynabeads Protein G kit (Invitrogen, Carlsbad, CA). Briefly, the anti-ATPB antibody was incubated with magnetic beads for 10 min at RT. The beads-antibody mixture was incubated with cell lysate for 2 hours at 4 °C, followed by elution with an elution buffer and LDS sample buffer for 10 min at 70 °C (Invitrogen, Carlsbad, CA). The beads were removed and immunoprecipitated proteins were analyzed using western blotting as previously reported (Zhang et al., 2016).

### Isolation of mitochondria and western blot analysis

Mitochondria were isolated from cells was performed following a published protocol (Clayton and Shadel, 2014). For western blotting, proteins were resolved by 10% SDS-PAGE and transferred onto a nitrocellulose membrane. After incubation with primary and secondary antibodies, membranes were scanned using the Odyssey CLx imaging system (LI-COR Biosciences, Lincoln, NE).

### Cell cycle analysis and mitochondrial ROS (mtROS) measurement in cultured cells

MDA-MB-231 cells were allowed to attach overnight. The medium was then switched to a glutamine-free DMEM medium containing 5% FBS and the test drug(s) for 48 hours. Cells were collected and stained with Propidium Iodide (PI) for 20-60 minutes, followed by an analysis using Attune Acoustic Focusing Cytometer (Applied Biosystems, Life Technologies). For mtROS measurement, MDA-MB-231 cells were cultured in the same conditions for 24 hours. Cells were stained with 5 μM MitoSOX for 20 minutes and analyzed using ZE5 Cell Analyzer (Bio-rad).

### Seahorse XF extracellular flux assay

To measure oxygen consumption rates, MDA-MB-231 cells were seeded were seeded in a concentration of 20,000 cells per cell into into XF96-well microplates (Seahorse Bioscience, 102416-100) and allowed to adhere for 8 hours. The cells were then changed into a glutamine-free DMEM media containing 5% FBS overnight followed by 4 hour drug treatment. They were subjected to the XF extracellular flux assay. Approximately 1 μM oligomycin, 1 μM FCCP and 0.5 μM Rotenone/Antimycin A mixture were spiked into cells at various time points. The oxygen consumption rate (OCR) was measured by the Seahorse XF extracellular flux assay. Eight replicates were run for each group.

### Statistical analysis

Statistical analyses were performed using GraphPad Prism 8 and in consultation with the University of Michigan’s Consulting for Statistics, Computing and Analytics Research (CSCAR) team. NaR cell proliferation index data were analyzed using pairwise Chi-square tests. All other data are shown as mean ± standard error (SEM). Statistical analyses were performed using unpaired two-tailed *t* test or ANOVA when appropriate. When comparing the means of multiple groups to a control group, Dunnett test or other tests were used to correct for multiple comparisons after ANOVA. Statistical significances from all tests were accepted at *p* < 0.05 or higher.

## Acknowledgments

We thank Dr. José M. Cuezva, Autonomous University of Madrid, Dr. Arohan R. Subramanya, University of Pittsburgh, Dr. Yonghua Sun, the Institute of Hydrobiology, Chinese Academy of Sciences; Dr. Heather Flanagan-Steet, Clemson University; and Dr. Yanzhuang Wang, University of Michigan, for providing reagents and fish lines. We are grateful to Dr. Chunyang Zhang for her advice on mitochondrial DNA quantification.

## Funding

This work was supported by NSF grant IOS-1557850 and NSF IOS-1755268 to CD. The funders had no role in study design, data collection and analysis, decision to publish, or preparation of the manuscript.

## Author contributions

Conceptualization: CD, YL, CL

Methodology: ZC, JL

Investigation: YL, CL, LR, VS, ZC, JG, CB

Visualization: YL

Supervision: CD

Writing—original draft: CD, YL

Writing—review & editing: YL, CL, LR, VS, ZC, JG, CB, JL, CD

## Competing interests

All authors declare they have no competing interests.

## Data and materials availability

Data generated or analyzed are included in the manuscript and supporting files.

## Supplementary Materials

**Supplementary Fig. 1.**
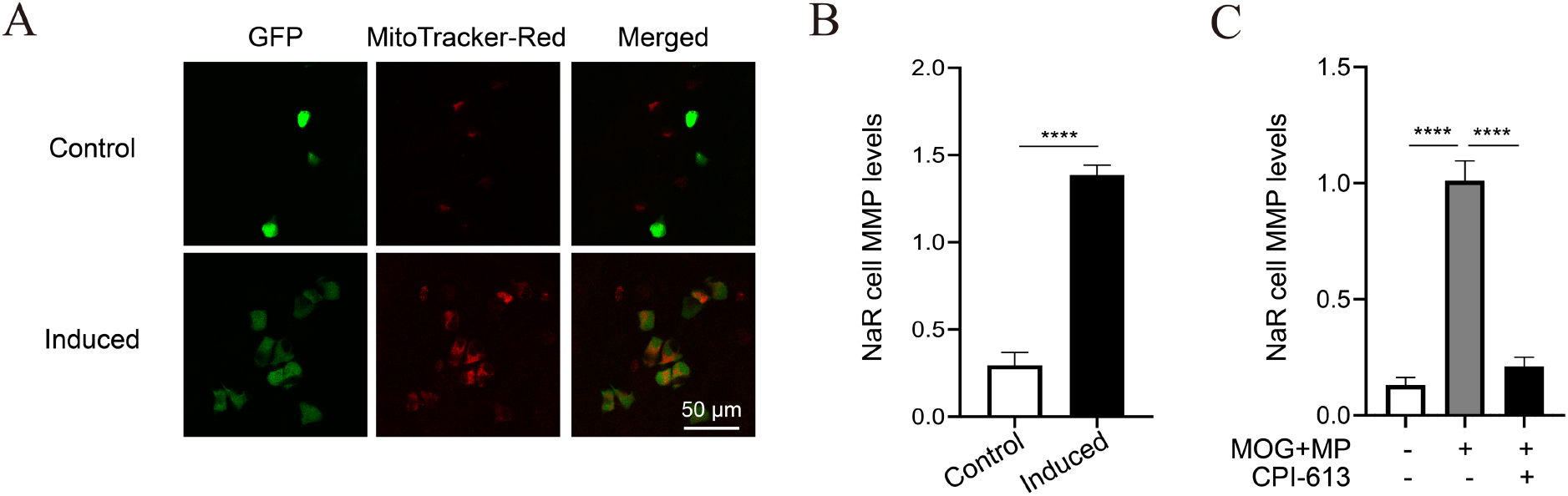
Increased mitochondrial membrane potential (MMP) in reactivated NaR cells. (A-B) *Tg(igfbp5a:GFP)* larvae (3 dpf) were transferred to the control or induction medium. Two days later, NaR cell MMP levels were measured by MitoTacker-Red staining. Representative images are shown in (A) and quantified results in (B). n= 216 to 672 cells from multiple fish. (C) MMP change is under the control of TCA cycle activity. *Tg(igfbp5a:GFP)* larvae were transferred to the control medium containing the indicated drugs at 3 dpf. Two days later, NaR cell MMP levels were measured after TMRM staining and shown. Dimethyl 2-oxoglutarate (MOG, 50 μM) and methyl pyruvate (MP, 50 μM); CPI-613, 3 μM.

**Supplementary Fig. 2.**
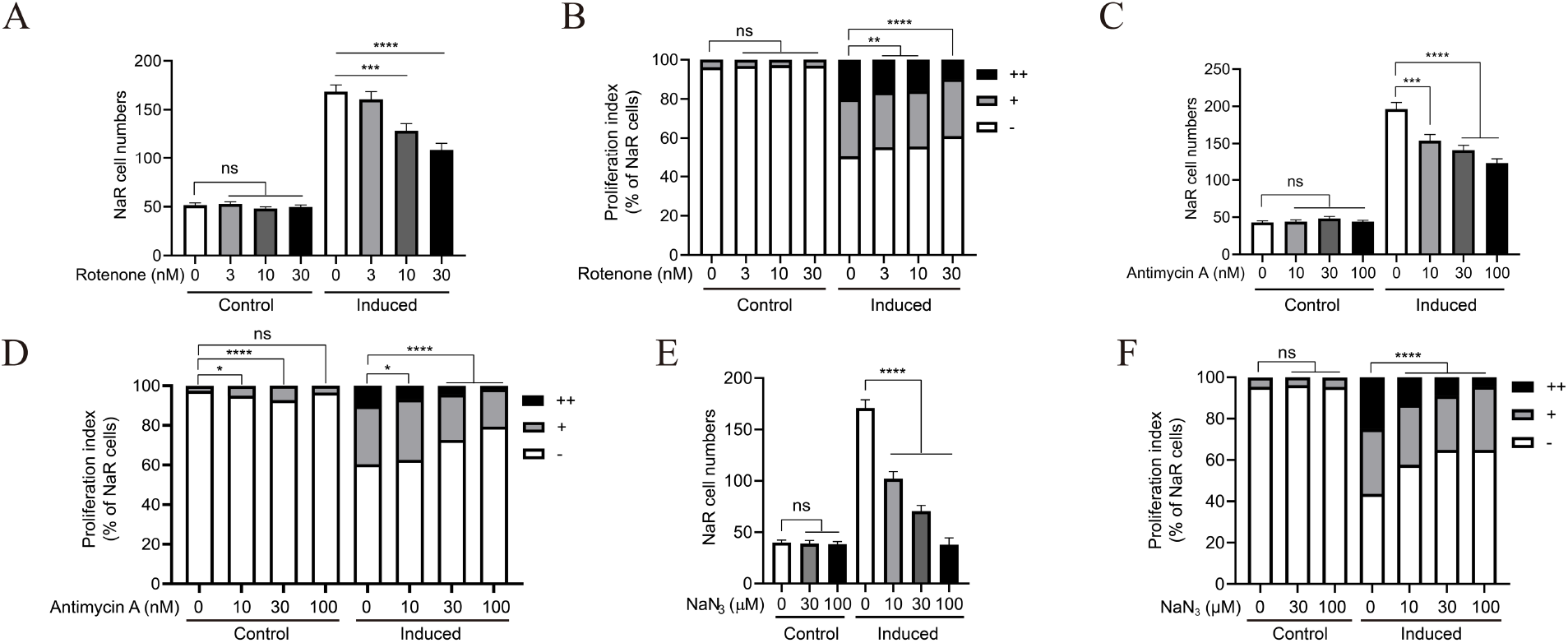
The ETC complex activity is required in NaR cell reactivation. (A-F) *Tg(igfbp5a:GFP)* fish (3 dpf) were transferred to the control or induction medium containing the indicated doses of ETC complex I inhibitor rotenone (A-B), ETC III inhibitor antimycin (C-D), and ETC IV inhibitor NaN3 (E-F), respectively. Two days later, NaR cell number and proliferation index were measured and shown. n = 9~33 fish/group.

**Supplementary Fig. 3.**
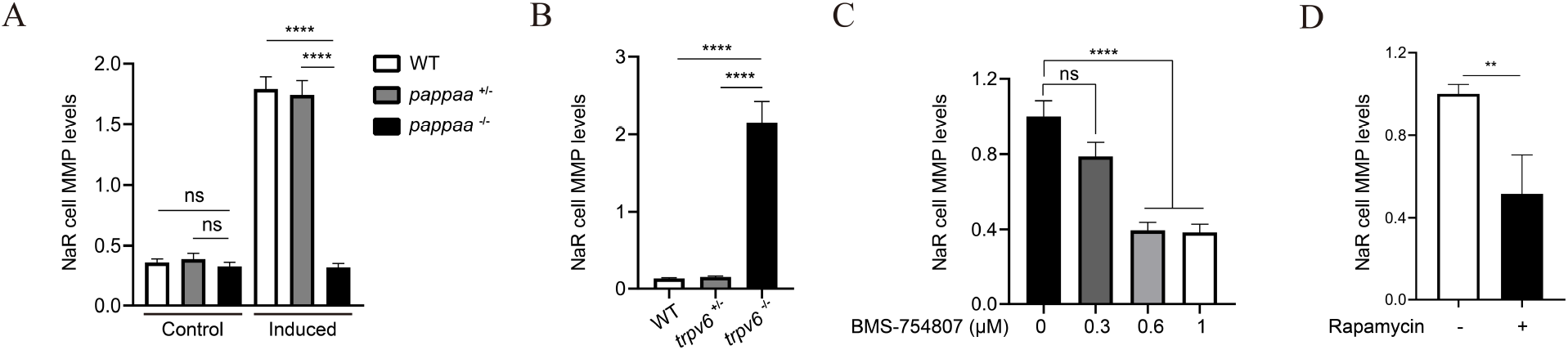
Mitochondrial membrane potential (MMP) increase is under the regulation of IGF and Tor signaling. (A) MMP levels in Igf signaling deficient NaR cells. Progeny of *pappaa^+/-^; Tg(igfbp5a:GFP)* intercrosses were raised to 3 dpf and transferred into the control or induction medium. NaR cell MMP levels were determined at 4 dpf followed by individual genotyping. n = 220 ~1242 cells from multiple fish. (B) MMP levels in Igf signaling active NaR cells. Progeny from *trpv6^+/-^; Tg(igfbp5a:GFP)* intercrosses were raised in the control medium and MMP was measured at 5 dpf. Each fish was genotyped afterwards. n= 222~2080 cells from multiple fish. (C-D) Effect of the IGF1 receptor inhibitor BMS-754807 and Tor inhibitor rapamycin. *Tg(igfbp5a:GFP)* larvae were transferred to the induction medium with or without the indicated concentrations of BMS-754807 (C) and rapamycin (5 μm) (D) for 1.5 hours and MMP was measured and shown. n = 88-347 cells from multiple fish.

**Supplementary Figure 4.**
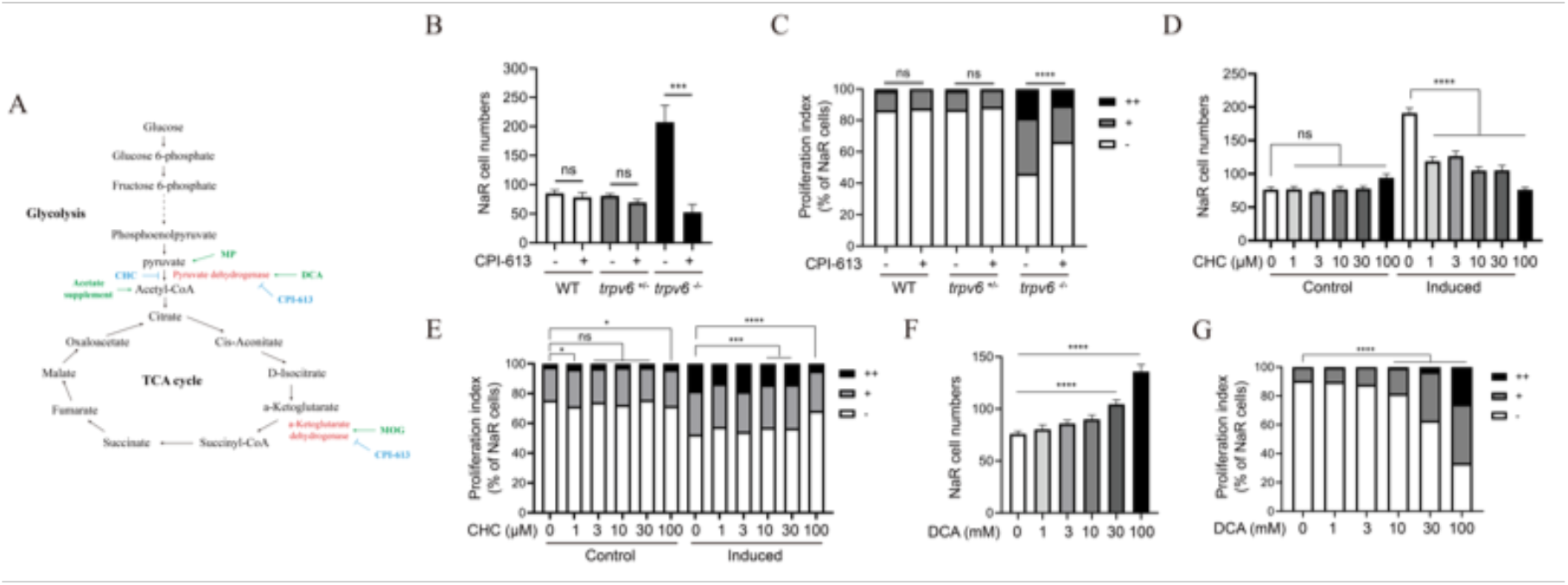
Elevated TCA cycle/OXOPO is required and sufficient in reactivating NaR cells. (A) Schematic diagram of glycolysis and TCA cycle and inhibitors/activator used in this study. MOG: Dimethyl α-ketoglutarate; MP: Methyl pyruvate; DCA: Dichloroacetate; CHC: α-cyano-4-hydroxycinnamate. (B-C) CPI-613 inhibits NaR cell proliferation in *trpv6^-/-^* fish. Progeny of *trpv6^+/-^; Tg(igfbp5a:GFP)* intercrosses were raised to 3 dpf and transferred into the control medium with or without 1 μM CPI-613. After two days, NaR cell number (B) and proliferation index (C) were measured and shown. n = 5~18 fish/group. (D-G) Effect of α-cyano-4-hydroxycinnamate (CHC) (D-E) and Dichloroacetate (DCA) (F-G). *Tg(igfbp5a:GFP)* embryos were raised and treated as described in Fig. 2 with the indicated doses of chemicals. NaR cell number (D, F) and proliferation index (E, G) were measured and shown. n= 24~34 fish/group.

**Supplementary Figure 5.**
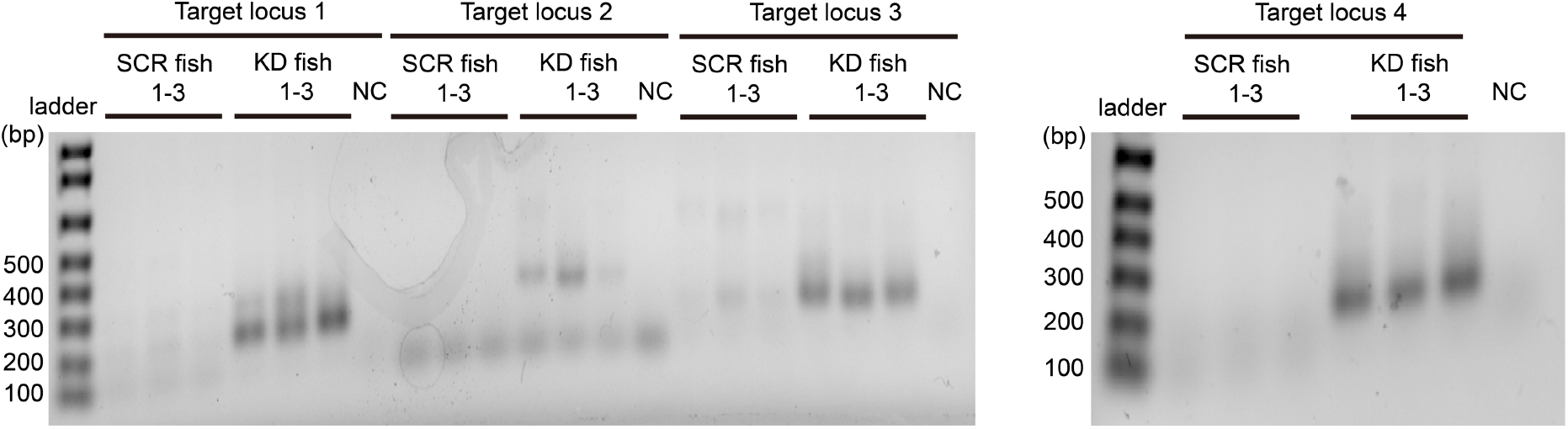
Validation of CRISPR/Cas9-mediated *atp5b* deletion. The deletion efficacy was confirmed by amplifying the 4 target loci in *atp5b* with head-loop primers following a recently published protocol (Kroll et al., 2021). SCR fish, fish injected with the scrambled gRNA; KD, fish injected with the indicated targeting gRNA; NC, negative control.

**Supplementary Figure 6.**
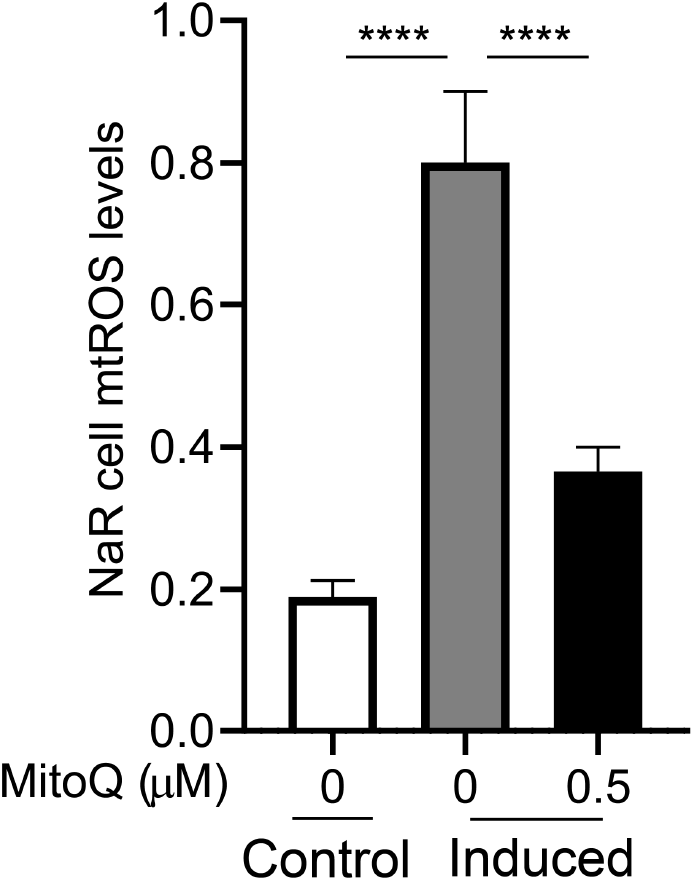
Validation of NaR cell mtROS staining. *Tg(igfbp5a:GFP)* larvae (3 dpf) were transferred to the control or induction medium in the presence or absence of the indicated doses of MitoQ. One day later, mtROS levels were measured and shown. n = 280-434 cells from multiple fish.

**Supplementary Figure 7.**
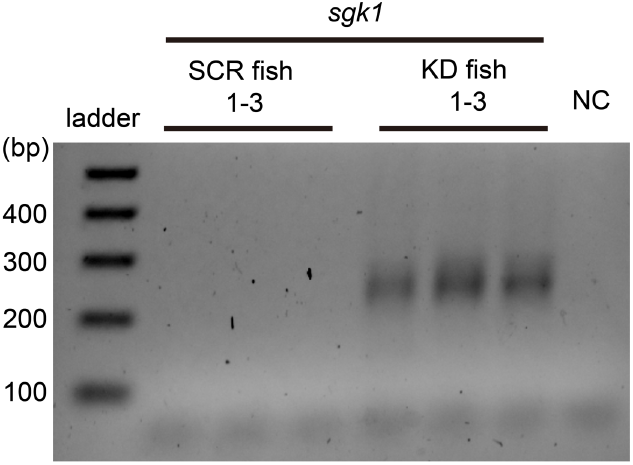
Validation of CRISPR/Cas9-mediated *sgk1* deletion. The knockdown efficacy was confirmed by amplifying a region covering all 4 target loci using head-loop PCR (Kroll et al., 2021). Arrowheads mark the 200 and 400 bp ladder band. scr, fish injected with the scrambled gRNA; sgk1, fish injected with sgk1 targeting gRNA; NC, negative control.

**Supplementary Figure 8.**
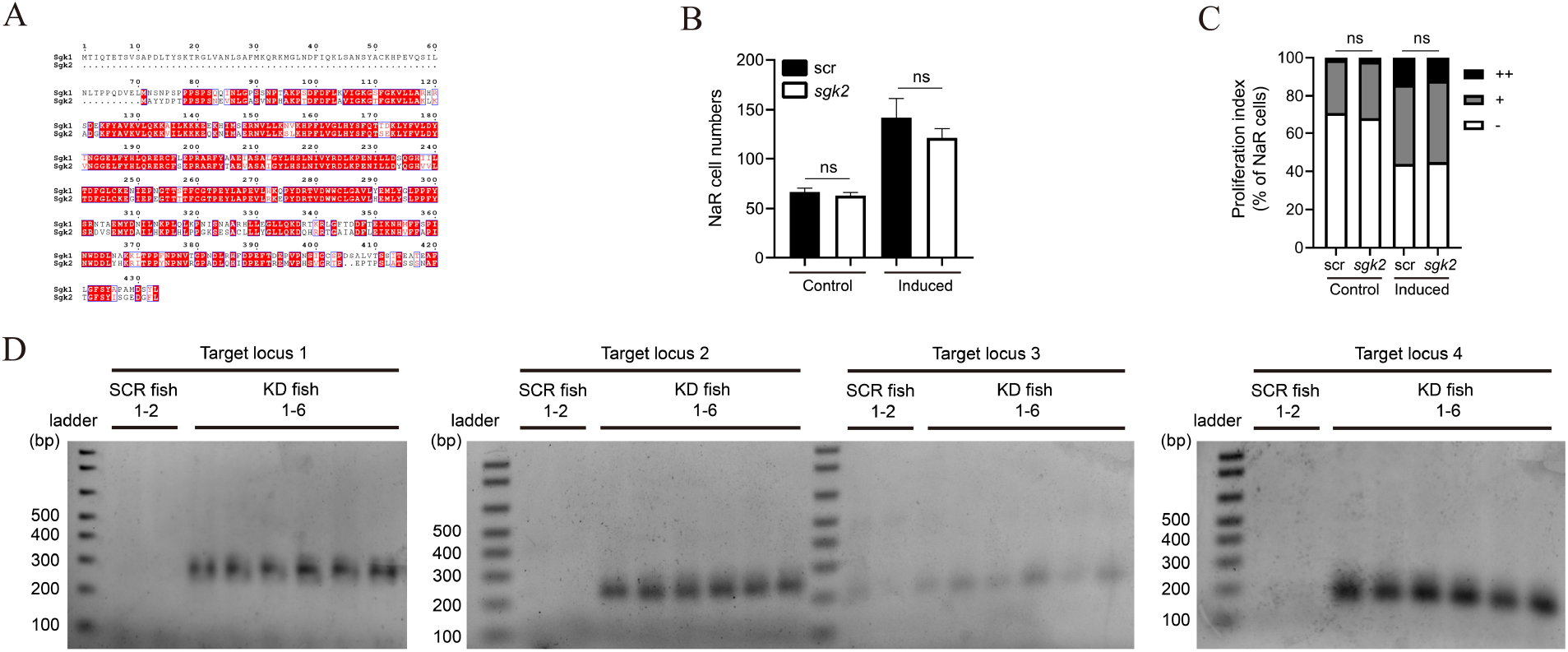
Sgk2 is not involved in NaR cell reactivation. (A) Amino acid sequence alignment of zebrafish Sgk1 and 2. Note the absence of the N-terminal sequence in Sgk2 where the mitochondrial targeting sequence is located. (B) CRISPR/Cas9-mediated *sgk2* deletion. *Tg(igfbp5a:GFP)* embryos were injected with *sgk2* targeting gRNAs and *Cas9* mRNA at the one-cell stage. They were raised to 3 dpf and transferred to the control or induction medium. Two days later, NaR cell number (B) and proliferation index (C) were measured and shown. n = 26-57 fish/group. (D) Validation of CRISPR/Cas9-mediated *sgk2* deletion. The knockdown efficacy was confirmed by amplifying the 4 target loci using head-loop PCR (Kroll et al., 2021). SCR, fish injected with the scrambled gRNA; KD fish, individual fish injected with the indicated *sgk2* targeting gRNAs.

**Supplementary Figure 9.**
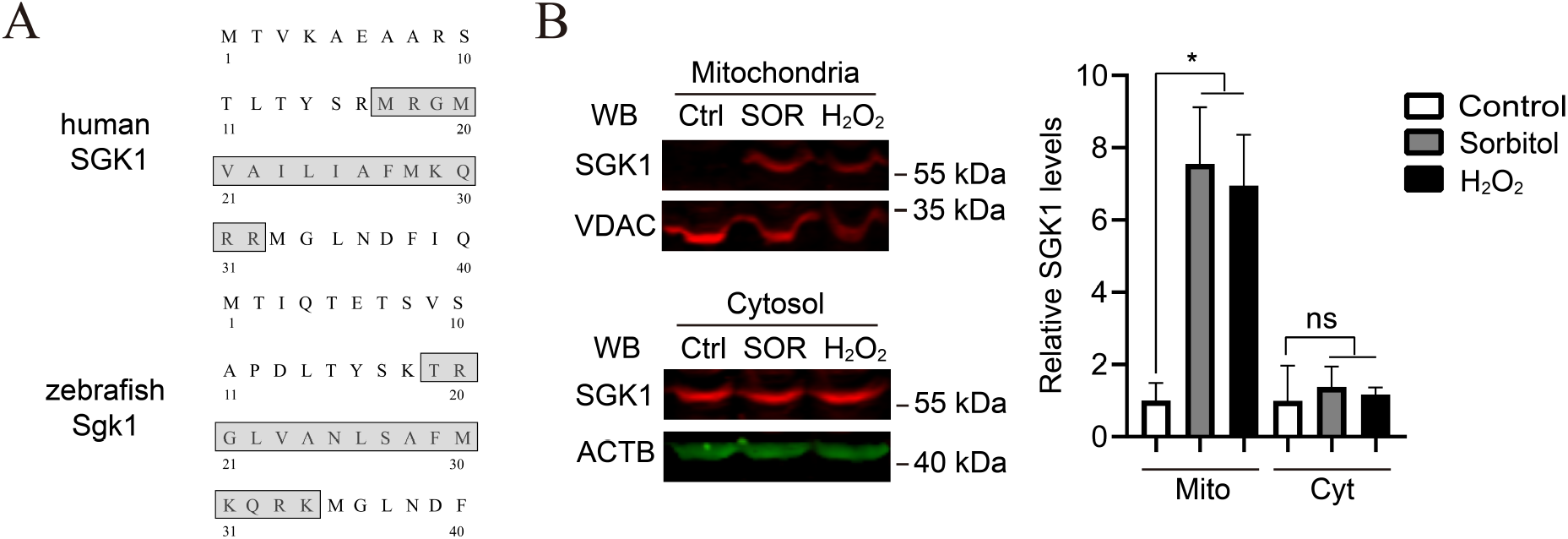
(A) Alignment of human SGK1 and zebrafish Sgk1 N-terminal sequences. The putative mitochondrial localization sequences are boxed. (B) ROS induces SGK1 expression in the mitochondria. MDA-MB-231 cells were treated with 0.3 M sorbitol for 18 hours or 250 μM H_2_O_2_ for 2 hours. Cells were lysed, fractioned, and examined by western blot using the indicated antibodies. Representative results were shown in the left panel and quantified results in the right panel. n = 3.

**Supplementary Figure 10.**
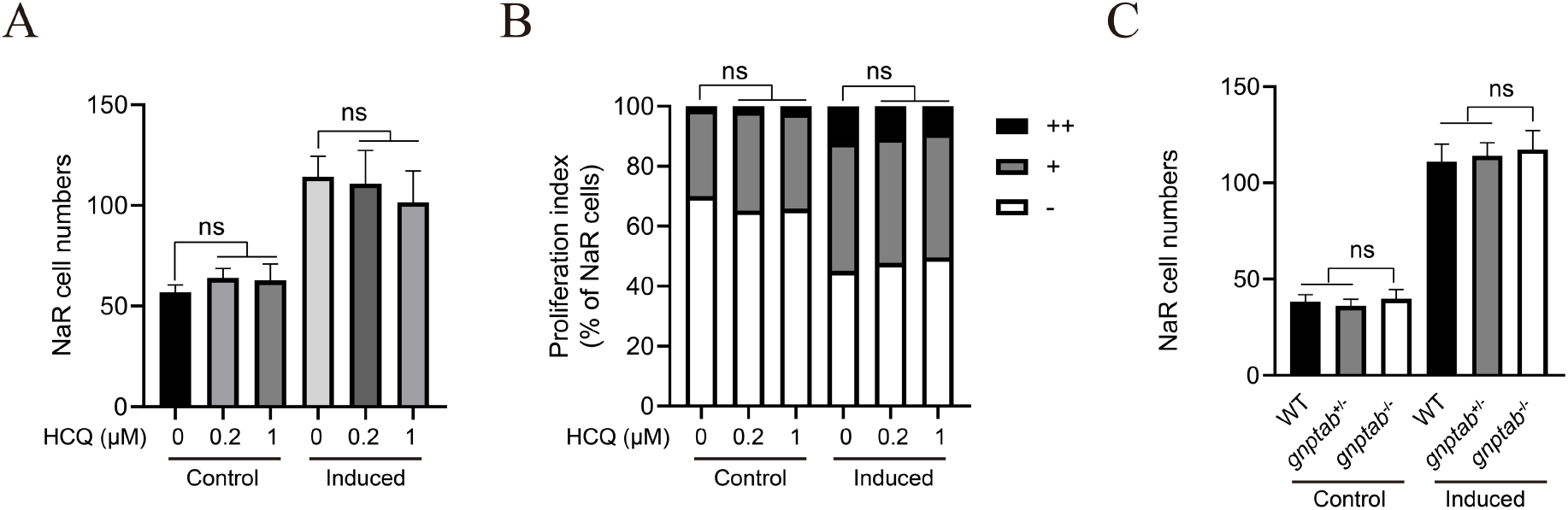
Autophagy is not involved in NaR cell reactivation. (A-B) Effect of hydroxychloroquine (HCQ). *Tg(igfbp5a:GFP)* larvae (3 dpf) were transferred into the control or induction medium in the presence or absence of the indicated doses of HCQ. Two days later, NaR cell number (A) and proliferation index (B) were measured and shown. n = 12-18 fish/group. (C) *Tg(igfbp5a:GFP)* larvae (3 dpf) of the indicated genotypes were into the control or induction medium in the presence or absence of the indicated doses of HCQ. Two days later, NaR cell number was measured and shown. n = 8-23 fish/group.

**Table S1.**
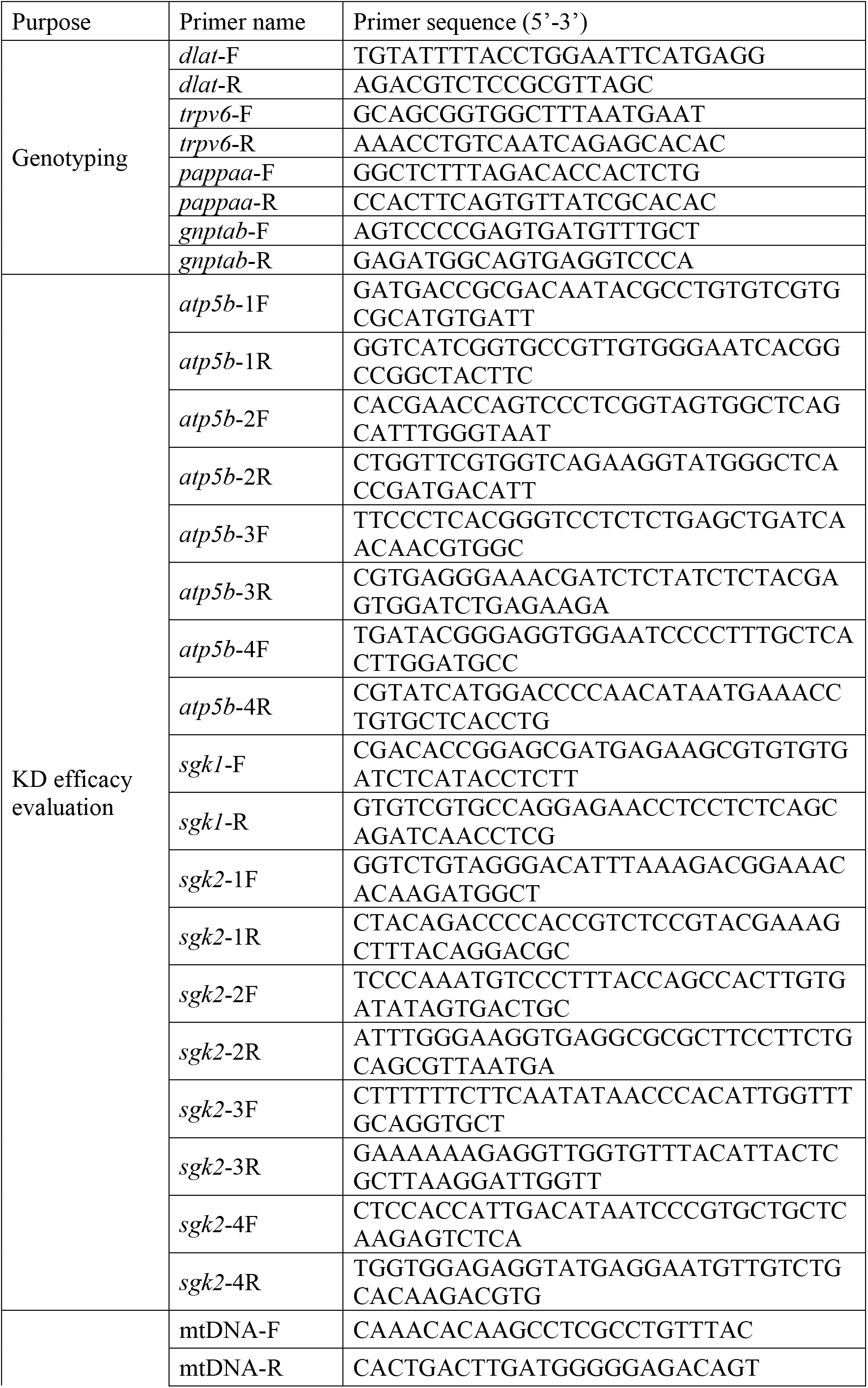
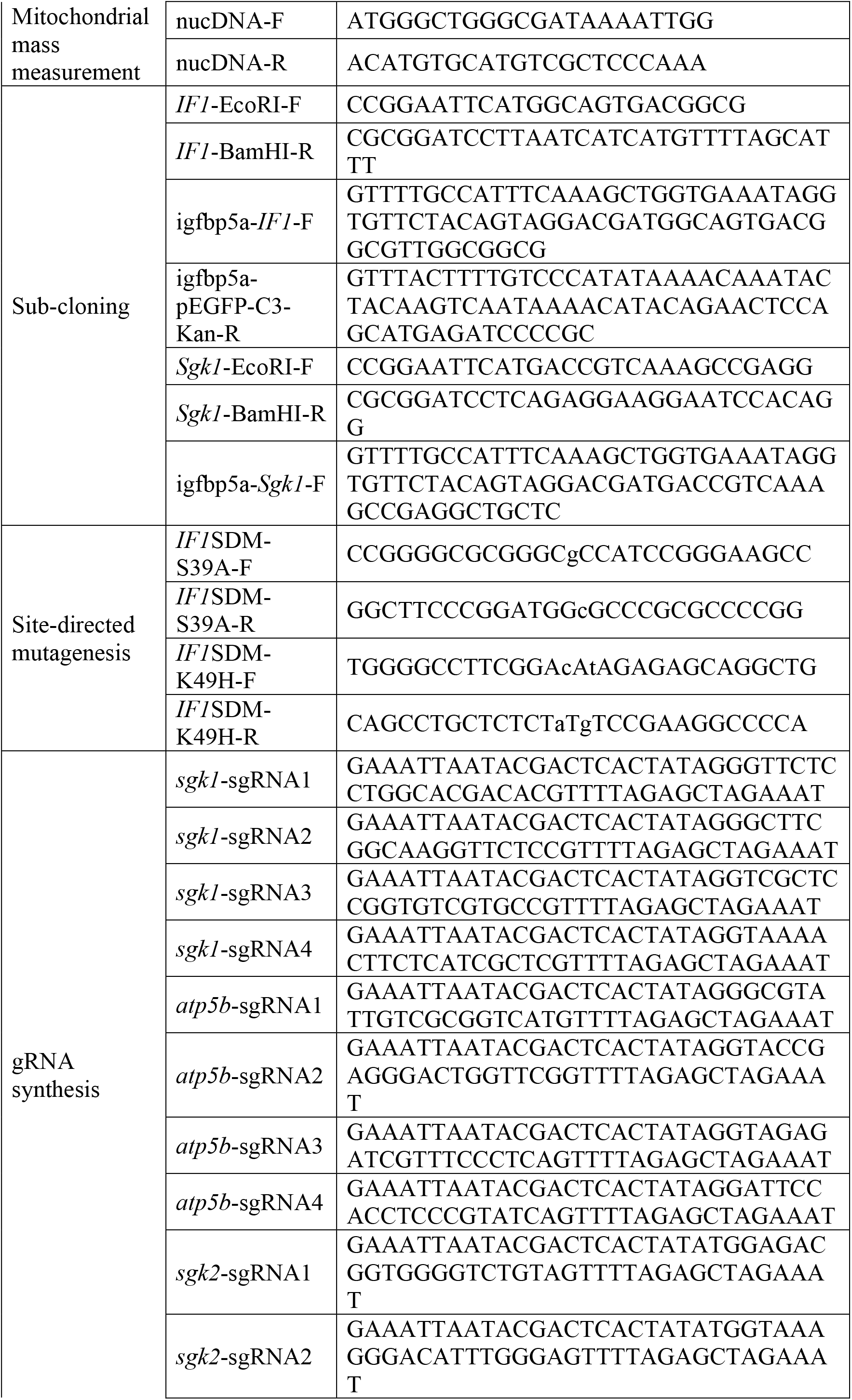
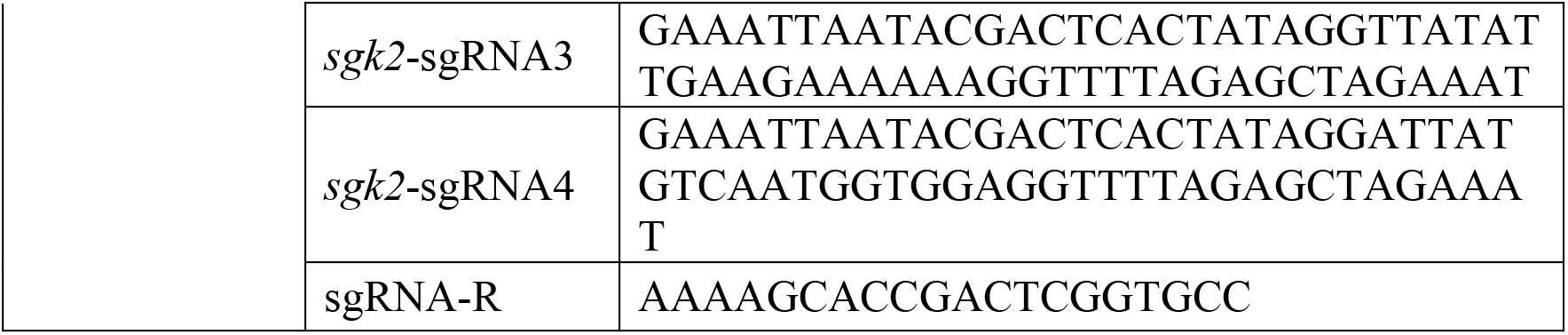
Primers used in this study.

## Notes

### Competing Interest Statement

The authors have declared no competing interest.

